# Suggesting disease associations for overlooked metabolites using literature from metabolic neighbours

**DOI:** 10.1101/2022.09.13.507596

**Authors:** M. Delmas, O. Filangi, C. Duperier, N. Paulhe, F. Vinson, P. Rodriguez-Mier, F. Giacomoni, F. Jourdan, C. Frainay

## Abstract

In human health research, metabolic signatures extracted from metabolomics data are a strong-added value for stratifying patients and identifying biomarkers. Nevertheless, one of the main challenges is to interpret and relate these lists of discriminant metabolites to pathological mechanisms. This task requires experts to combine their knowledge with information extracted from databases and the scientific literature.

However, we show that a large fraction of metabolites are rarely or never mentioned in the literature. Consequently, these overlooked metabolites are often set aside and the interpretation of metabolic signatures is restricted to a subset of the significant metabolites. To suggest potential pathological phenotypes related to these understudied metabolites, we extend the ‘guilt by association’ principle to literature information by using a Bayesian framework. With this approach, we suggest more than 35,000 associations between 1,047 overlooked metabolites and 3,288 diseases (or disease families). All these newly inferred associations are freely available on the FORUM ftp server (See information at https://github.com/eMetaboHUB/Forum-LiteraturePropagation.).

## 1 Introduction

Omics experiments have become widespread in biomedical research, and are frequently used to study pathologies at the genome, transcriptome, proteome and metabolome levels. The subsequent discriminant analysis leads to a set (a signature) of genes, proteins or metabolites, reflecting alterations of the phenotype at different levels of post-genomic processes. The interpretation of these signatures requires gathering knowledge about each of its elements from the scientific literature and dedicated databases (DisGeNET[1], Uniprot[2], HMDB[3], CTD[4], MarkerDB[5], FORUM[6]). However, despite its exponential growth[7], the scientific literature suffers from an imbalanced knowledge distribution. This topic has received much attention for genes and proteins, showing a highly skewed distribution of the number of articles mentioning each entity. Consequently, this strong imbalance has an impact on the quantity and quality of gene annotations in databases[8, 9, 10, 11, 12]. Indeed, what is known as *the Matthew effect* [13], which refers to the saying “the rich get richer”, is particularly valid in scientific communications. For instance, as reported in [9]: “*more than 75% of protein research still focuses on the 10% of proteins that were known before the genome was mapped* “ and as reported in [12] “*all genes that had been reported upon by 1991 (corresponding to 16% of all genes) account for 49% of the literature of the year 2015*.”.

Metablomics emerged later than its omics siblings, transcriptomics and proteomics, and has, like them, benefited from technological advancements, such as NMR and mass spectrometry. While we are getting closer to a complete reconstruction of the human genome[14], our knowledge of the metabolome, i.e. the set of metabolites present in a biological system[15], is still limited by technical constraints. Among them, the main limitations are the identification of unknown metabolites and the sometimes inaccurate identification of known ones [16, 17]. For instance, only a small fraction (< 20%) of metabolic spectra can be correctly annotated [18, 19] in an untargeted metabolic analysis. This disparity is also reflected in the distribution of the number of articles mentioning each compound present in the PubChem Database. While only a small fraction of them are mentioned in thousands of articles, the majority remains rarely or never mentioned [20]. This imbalance has consequences for the interpretation of the signatures, which can rely solely on a subset of its members that are sufficiently covered to provide insights. In Human health research, it is therefore critical to bring knowledge to these understudied compounds, by suggesting diseases that could be linked to them.

A metabolite is suspected to be impacted or involved in a particular disease through metabolism when an imbalance in its abundance has been observed in comparison to control cases. Moreover, metabolites are linked to each other by biochemical reactions, and therefore their abundances are also interdependent.

Among other factors, the abundance of a compound can depend on the concentration of its precursors and, in turn, can also influence the rate of production of other compounds. Following the well known ‘guilt by association’ hypothesis, we assume that: if a metabolite has been linked to a particular disease due to an imbalance in its abundance, metabolites that are connected to it by biochemical reactions, i.e. its metabolic neighbourhood, can also be suspected of being linked to this disease. Metabolic networks[21], built originally for modelling purposes, describe those substrate-product relations between compounds and thus provide a suitable support to extend these suspicions to metabolic neighbours. For Human, the reconstruction of the metabolic network (Human1 v1.7 [22]) contains 13,082 reactions and 8,378 metabolites. In other omics fields, network-based strategies following “guilt-by-association” principle have been applied to build several recommendation systems proposing new genes or proteins that could be related to a given disease from a list of known genes/proteins [23, 24, 25]. We also developed a similar approach for metabolic signatures using random walks in metabolic networks [26].

If a compound is rarely or never mentioned, we hypothetize that the literature in its surrounding neighbourhood may provide *a priori* knowledge on its biomedical context. To combine both this *a priori* and the available literature of the compound (if any) in the suggestions, we propose a method based on the Bayesian framework. The method returns several predictors to evaluate whether a significant proportion of the articles mentioning a metabolite would also mention a disease. In addition, several indicators can be used to highlight the most influential metabolic neighbours in the suggestions.

All the required data were extracted from the FORUM Knowledge Graph (KG)[6]. FORUM contains significant associations between PubChem chemical compounds and MeSH biomedical descriptors based on their co-mention frequency in PubMed articles. We evaluated our hypothesis by testing whether significant associations between metabolites and diseases could be retrieved solely on the basis of the literature of their neighbours. We illustrate the behaviour of the method in two scenarios: a metabolite without available literature for which the prior is the only source of information (Hydroxytyrosol) and a rarely mentioned metabolite (5*α*-androstane-3,17-dione with 82 articles). Using this approach on human metabolic network, we suggested more than 35,000 new relations between overlooked metabolites and diseases (and disease families). The code and the data needed to reproduce the results are available at https://github.com/eMetaboHUB/Forum-LiteraturePropagation.

## 2 Method Summary

The core of the method is the construction of a prior distribution on the probability that an article mentioning a metabolite would also mention a particular disease. This distribution is estimated from the literature of its metabolic neighbourhood. The metabolic neighbourhood of a compound consists of the metabolites that can be reached through a sequence of biochemical reactions. It is defined from the Human1 metabolic network[22], which was pruned from spurious connections using an atom-mapping procedure[26] (see S1.1). In the following description of the method and subsequent analyses, overlooked metabolites will be divided into two categories: those without literature (1) and those that are rarely mentioned (2).

The Figure 1 summarizes all the steps in the proposed method. Figure 1.A introduces the example of a relation between an overlooked metabolite *A* and a disease. The prior distribution on the probability that an article mentioning *A*, would also mention the disease, is built from a mixture of the literature of its close neighbourhood in the metabolic network. The weight of the component of these metabolites in the mixture, depends both on their distance to *A* and their amount of literature (see details in 7.2). We impose that a metabolite can’t influence its own prior or the prior of far distant metabolites. As an illustration, *B* shares a quantity *t*_*B,A*_ of its literature to build the prior of *A*, but doesn’t influence its own prior, as well as the prior of *Z* (Cf. Figure 1.B). The weight of *B* in the prior of *A* is then estimated as the amount of literature it had shared with *A*, relative to the other neighbours *C, D, F* (See Figure 1.C). We refer to *B, C, D* and *F* as the *contributors* to the prior of *A*. By analogy, it is as if each metabolite spreads its literature in the metabolic network, and the prior of *A* was built from the articles it had received from its contributors.

**Figure 1:**
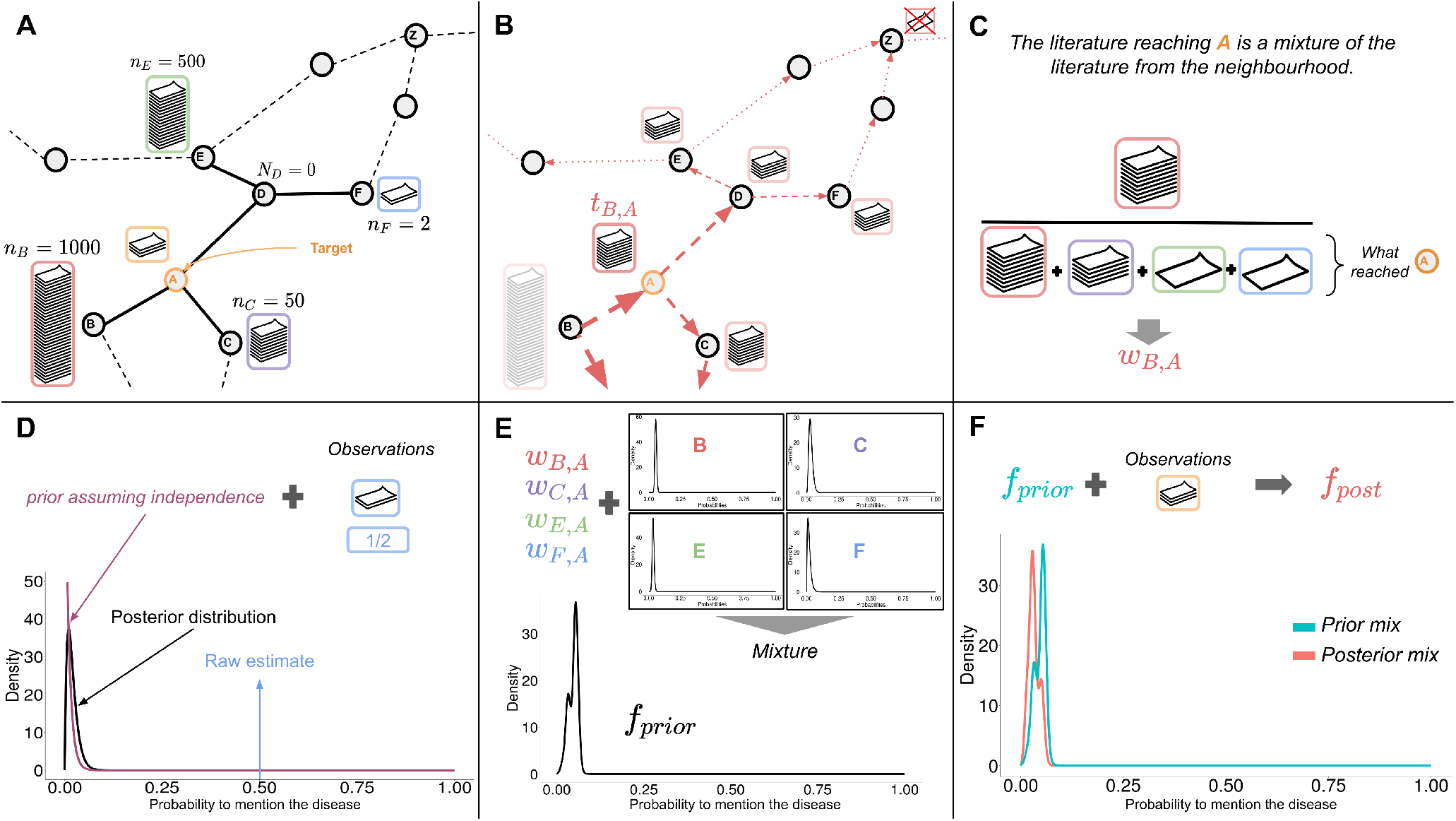
A step by step description of the proposed method. Compound A has 0 < *n*_*A*_ ≤ 100 articles, with some co-occurrence with the disease of interest (0 ≤ *y*_*A*_ ≤*n*_*A*_). In the blocks **A** and **B**, the nodes represent metabolites and the edges substrate/product relationships in the metabolic network. Dashed lines indicate more distant connections.

In Figure 1.D, the contributor *F* is also an overlooked metabolite with only 2 annotated articles, including one mentioning the disease. This lack of literature may lead to a less reliable contribution. To avoid this issue, an initial shrinkage procedure is applied to all contributors. The probability distribution that one of its articles mentions the disease is readjusted toward the overall probability of mentioning the disease (see details in 7.3).

Then, we build the prior distribution for *A*, by mixing the probability distributions of each contributor (see Figure 1.E) according to their weights estimated in the previous step (Figure 1.C). The prior mixture distribution is denoted by *f*_*prior*_. The constructed prior distribution for *A* represents the probability distribution that an article from one of its contributors would mention the disease. In the scenario where *A* has no literature (1), the predictions will be based solely on *f*_*prior*_.

However if *A* is mentioned in few articles (2), we compute the posterior distribution, thus updating the weights and distributions of each contributor in the mixture (Figure 1.E). The posterior mixture distribution is denoted by *f*_*post*_.

From the mixture distribution, two predictors are estimated: *LogOdds* and *Log*_2_*FC. LogOdds* expresses the ratio between the probability of the disease being mentioned more frequently than expected in the literature of the compound, rather than less frequently. *Log*_2_*FC* expresses the change between the average probability of mentioning the disease in the mixture distribution, compared to the expected probability in the whole literature. In summary, both should be considered jointly in the predictions: *LogOdds* as a measure of significance and *Log*_2_*FC* as a measure of effect size. In (2), to get an intuition about the belief of the neighbourhood only, we also return similar indicators estimated from *f*_*prior*_: *priorLogOdds* and *priorLog*_2_*FC* (see details in 7.4 and 7.5). Finally, several diagnostic values such as *Entropy* allow to assess the composition of the built prior (See S1.3). *Entropy* evaluates the good balance of contributions in the prior. The more metabolites contribute to the mixture and the more their weights are uniformly distributed, the higher the entropy.

## 3 Results

### 3.1 Unbalanced distribution of the literature related to chemical compounds

The FORUM KG links PubChem compounds to the PubMed articles that mention them. Among the 103 million PubChem compounds in FORUM, only 376,508 are mentioned in PubMed articles, representing a coverage lower than 0.4%. For these mentioned compounds, the distribution of the literature is highly skewed (Figure 2.A). The top 1% of the most mentioned compounds (red area) concentrates 80% of the links between PubChem compounds and PubMed articles. Similarly, the blue area indicates that 63% of compounds (218,291) have only one article mentioning them, which, to give a point of comparison, is less than the literature associated with glucose: 278,277 distinct articles.

**Figure 2:**
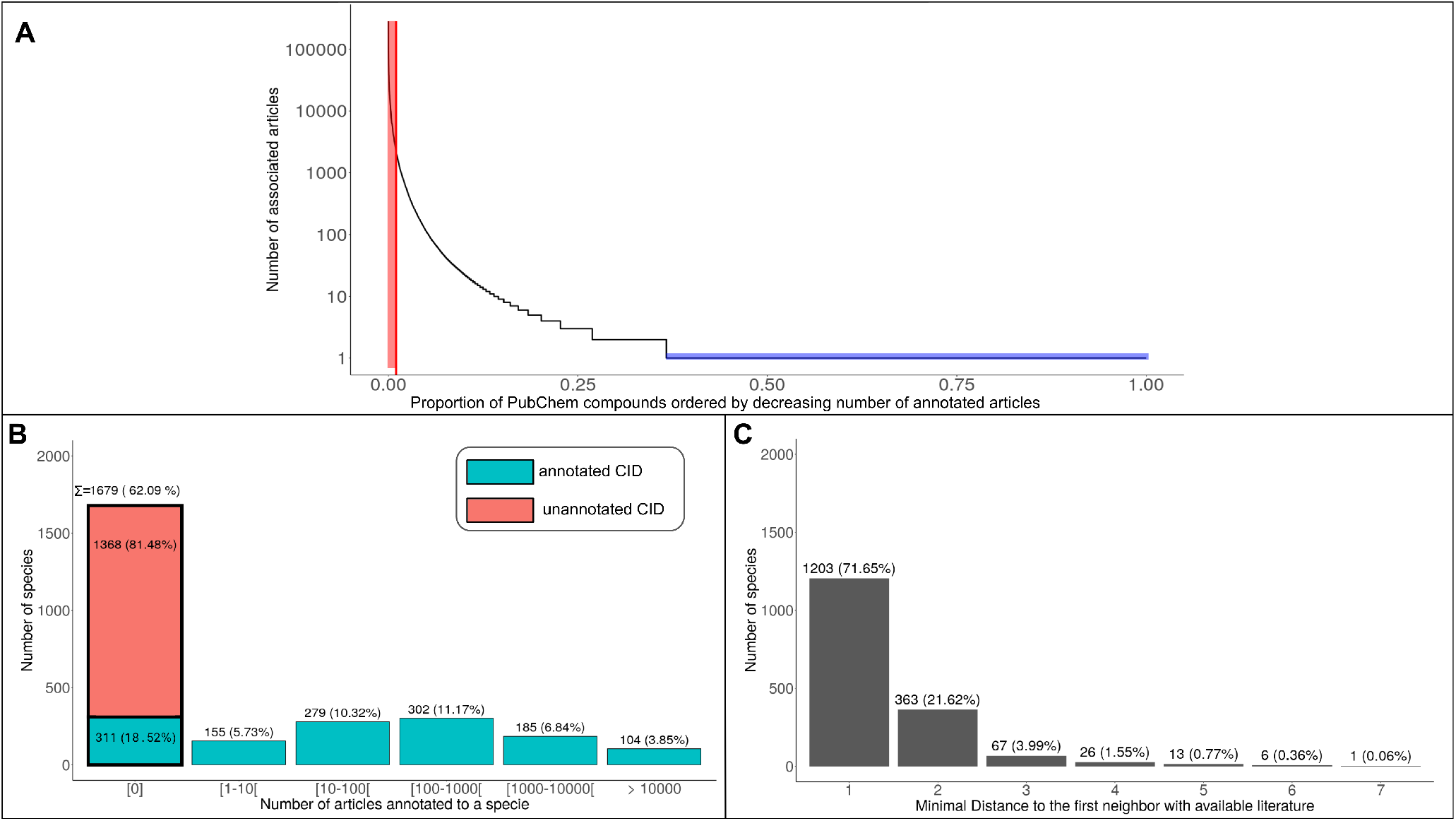
**A:** Distribution of the number of annotated articles (expressed in log-scale) for PubChem compounds that have at least one article in FORUM, in descending order. The red area represents the proportion of the most mentioned compounds required to attain 80% of the total number of annotations, while the blue area represents the fraction of compounds with only one annotated article. **B**: Distribution of the number of annotated articles per metabolites, organised by bins, in the carbon skeleton graph of Human1. The first bar represents the metabolites without literature. Among them, 81.5% don’t have annotated PubChem identifiers, making it impossible to link them to PubMed articles with FORUM. The remaining 16.5% have annotated PubChem identifiers, but no articles were found mentioning them. In total, there are 1336 articles with an available PubChem identifier. **C**: Distribution of the shortest distance to the first neighbour in the metabolic network with at least one annotated article, for the metabolites without literature in the network (bold bar of **B**). The distances were computed with the Dijkstra algorithm.

Considering only metabolites, Figure 2.B presents the distribution of the number of articles mentioning the 2704 metabolites, conserved in the pruned version of the Human1 metabolic network. Because of the skewed distribution of the literature and the lack of external identifiers, 62.09% of the metabolites in the metabolic network have no annotated articles. Nevertheless, almost 72% of them have at least one direct neighbour in the metabolic network with available literature (See Figure 2.C). Moreover, by considering the close neighbourhood (paths up to three reactions), almost all the metabolites (≈ 97.26%) without initial literature can reach a described neighbour, showing the availability of nearby literature to build a prior.

### 3.2 Evaluation of the prior computation

The critical step in the proposed method is the construction of a relevant prior. While its influence on the results will decrease as the size of the literature of the targeted compound increases, it will mainly drive the predictions for the rarely mentioned compounds we are interested in [27].

The relevance of the prior was evaluated by testing whether significant associations with diseases, could be retrieved using only the literature from the metabolic neighbourhood of the metabolite. The validation dataset includes 10,000 significant relations between metabolites and disease-related MeSH extracted from the FORUM KG, and 10,000 random metabolite-MeSH pairs to serve as negative examples. The method is evaluated by considering either the direct or a larger neighbourhood (metabolites that can be reached through a path of two or more reactions). In the method, the considered neighbourhood is controlled by the parameter *α* (see details in 7.2 and S4.1) and is set to *α* = 0 for the direct neighbourhood and *α* = 0.4 for a larger one.

We decided to compare the proposed method against two different baselines (more details in S4.2). Baseline-Freq is the most naive approach in which the predictions are solely based on the overall probability of mentioning the disease, such that a metabolite is more likely to be related to frequently mentioned diseases in the literature. Hence, Baseline-Freq ignores the network information (metabolic neighbourhood). On the contrary, the predictions with Baseline-DN are based on the average probability of mentioning the disease in the direct neighbourhood, thus closer to the proposed approach. It is worth noting that, if all direct neighbours have relatively the same amount of annotated articles and are well covered (negligible shrinkage), the method parameterized with *α* = 0 behaves like the simple Baseline-DN for metabolites without literature. We used *Log*_2_*FC* as predictor for the proposed method in Figure 3.

**Figure 3:**
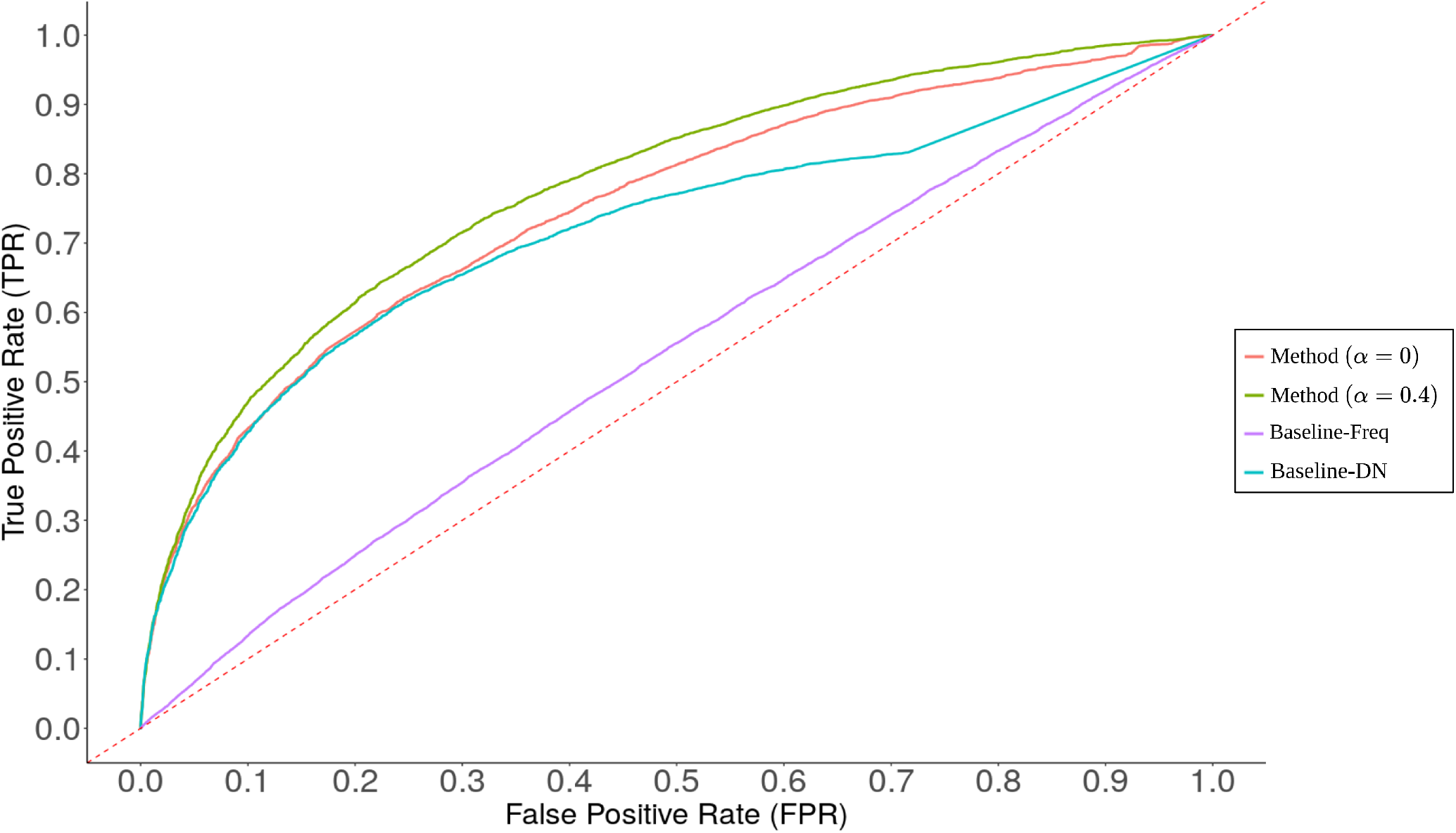
Receiver operating characteristic (ROC) of the method considering only the direct neighbourhood (*α* = 0) or a larger (*α* = 0.4) and two different baselines. For Baseline-Freq the predictions are only based on the overall probability of mentioning the disease in the literature. For Baseline-DN the predictions are based on the ratio between the average probability of mentioning the disease in the direct neighbourhood and its overall probability. Respective AUC (Area Under the Curve) for Method (*α* = 0), Method (*α* = 0.4), Baseline-DN and Baseline-Freq are: 0.75, 0.78, 0.72 and 0.54.

All tested approaches outperform Baseline-Freq, showing the benefit of examining the neighbouring literature. When considering the direct neighbourhood (method with *α* = 0), the method is more efficient than Baseline-DN. However, as previously shown in Figure 2.C, the direct neighbourhood cannot bring information for more than 28% of metabolites without literature. Therefore, considering a larger neighbourhood can be essential for some overlooked metabolites, and the approach achieves solid performances (AUC=0.78) on the validation dataset with *α* = 0.4. Applying a threshold on *Log*_2_*FC* > 1 results in a TPR=0.35 and a FPR=0.05. Using *LogOdds* as predictor, the method achieved slightly lower performances (AUC=0.76), with a TPR=0.22 and a FPR=0.04 when applying a threshold on *LogOdds* > 2. Beyond the validation, *LogOdds* is more robust to outlier contributions than *Log*_2_*FC* and when examining predictions, they should be considered together as complementary indicators of significance and effect size. These results suggest that the prior built from the neighbouring literature alone, holds relevant information about the biomedical context of metabolites and could be efficient to drive predictions for rarely mentioned compounds. To evaluate the performance of predictions based on the posterior distribution, a complementary analysis is provided in S4.3. Finally, as mentioned in the Method summary, the metabolic network was pruned from spurious connections using an atom-mapping procedure (see S1.1). The benefit of this procedure on the predictions is evaluated in S4.4.

### 3.3 Suggesting relations with diseases for overlooked metabolites

In the FORUM KG, 80% of the significant associations with biomedical concepts are observed for the 20% of compounds with more than 100 annotated articles. This manifestation of the Pareto principle[28] reflects the need for additional knowledge for compounds that are less frequently mentioned. Therefore in this analysis, we applied the proposed method on all metabolites in the human metabolic network with less than 100 annotated articles (see Table 1). According to the experiments on the validation dataset (See. 3.2), we applied a threshold on *LogOdds* > 2 and *Log*_2_*FC* > 1. We also retained predictions based on well-balanced contributions from the neighbourhood by filtering on the diagnostic indicator *Entropy* > 1 (See details in Method and S1.3).

**Table 1:**
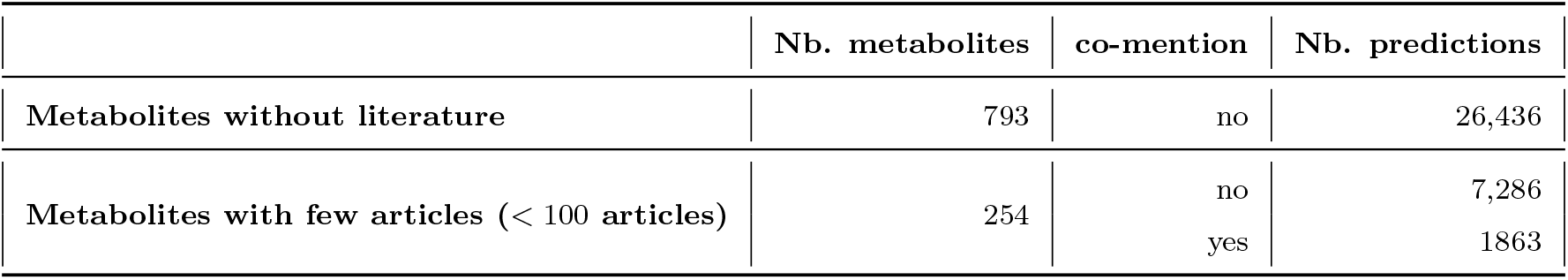
Summary table of the number of disease-related MeSH predicted for metabolites in the network with less than 100 annotated articles. The results are separated between the two major scenarios: (1) Metabolites without literature and (2) metabolites poorly described in the literature (< 100 articles). In the second case, results are also arranged according to whether the metabolite already co-mentions the MeSH (co-mention column). Only predictions with *LogOdds* > 2, *Log*_2_*FC* > 1 and *Entropy* > 1 are considered. For the 1863 predictions where the metabolite co-mentions the MeSH, 938 (≈ 50%) are also retrieved using a right-tailed Fisher exact test (BH correction and *q*.*value* < 0.05). Only 793 metabolites among the 1679 without literature and 254 among those with literature have significant results according to the used thresholds.

1863 predictions correspond to relations that are not novel, since they are already supported by one or several publications in the literature (`co-mention:yes` in Table 1). However, by re-evaluating these predictions using a right-tailed Fisher exact Test (BH correction and selecting those with *q*.*value* < = 0.05), we found that ≈ 50% of them (925) would not have been found significant. These relations are still weakly supported, nevertheless, our method showed that they are consistent with the neighbourhood. 7,286 novel relations have also been suggested with disease-related MeSH, without having already been mentioned in their literature (`co-mention:no`). Finally, for 793 metabolites without literature, 26,436 relations have been suggested only by exploiting the neighbourhood literature. All the results are available on the FORUM ftp server (See https://github.com/eMetaboHUB/Forum-LiteraturePropagation.), filling a gap when it comes to the interpretation of signatures with these understudied metabolites.

### 3.4 Case study

In this section, we will describe the behaviour and benefits of the method through two test cases. As mentioned in 2, Hydroxytyrosol is an example of a metabolite without literature (1) and 5alpha-androstane-3,17-dione of a metabolite with only a few annotated articles (2) and with a weakly supported association.

#### 3.4.1 Hydroxytyrosol and its potential link with Parkinson’s disease

Hydroxytyrosol is a metabolite which is known for its antioxidant properties [29] and mentioned by 856 publications in FORUM. However, its literature will only serve as ground truth, and Hydroxytyrosol will be considered as a metabolite without literature in this analysis. Consequently, the predictions are solely derived from the neighbouring literature (*f*_*prior*_). The top 10 predictions ranked by *LogOdds* are presented in Supplementary Table S1. Parkinson’s disease is the most suggested disease, followed by broader descriptors also related to neurodegenerative disorders. This suggestion is mainly driven by the literature of close metabolic neighbours: dopamine and 3,4-dihydroxyphenylacetate (Figure 4). Both compounds’ literature frequently mention Parkinson’s Disease (Table S2) suggesting that Hydroxytyrosol may also be related to this disease. Other contributors such as 3.4-dihydroxyphenylacetaldehyde or homovanillate also seem to be related to the pathology but only contribute ≈ 5% to the prior as they are more distant neighbours or have less literature. In the actual literature of Hydroxytyrosol, 2 articles[30, 31] explicitly discuss its therapeutic properties on Parkinson’s disease.

**Figure 4:**
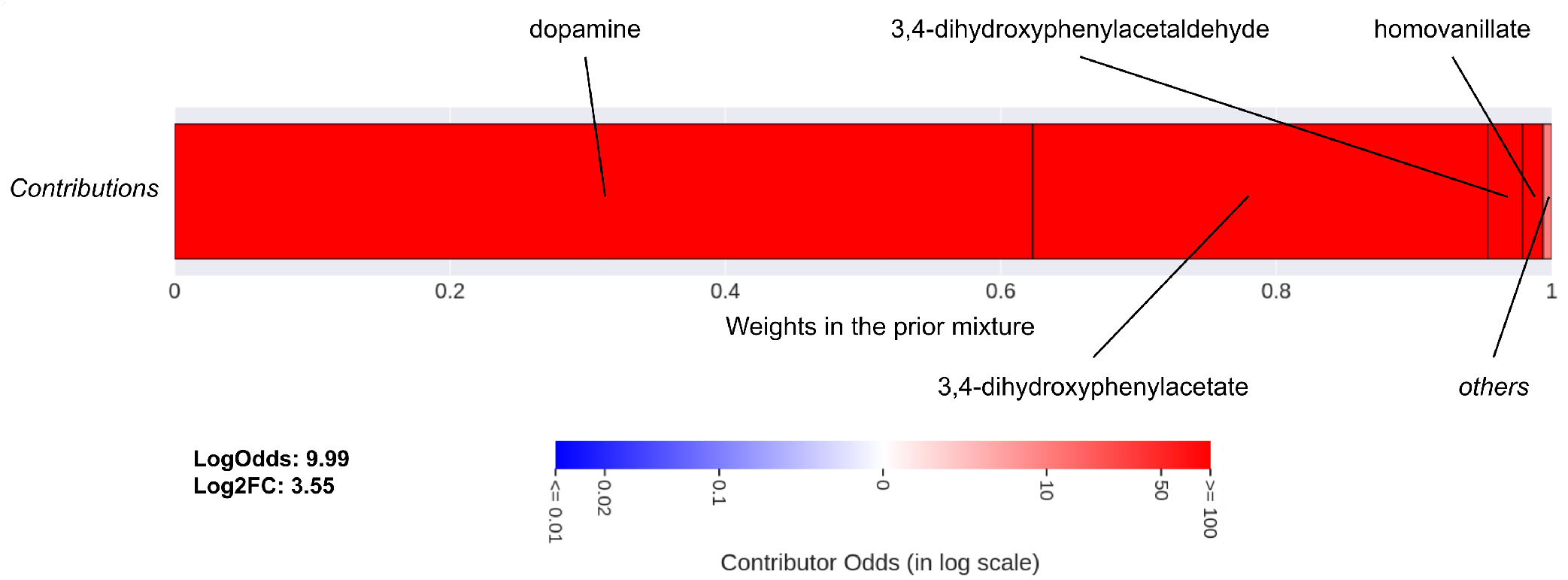
Profile of the contributors for the association between Hydroxytyrosol and Parkinson’s Disease. This shows the repartition of the literature received by Hydroxytyrosol from its neighbourhood to build its *prior*. Contributors are organised in blocks by increasing weights in the prior mixture (*w*_*i,k*_), from left to right. The weights also give the width of the block. The colour of each block associated with a contributor depends on its individual *LogOdds*, from blue to red, for *negative* (less likely) to *positive* (more likely) contributions respectively. Weights and *LogOdds* are also detailed in table S2.

#### 3.4.2 Highlighting the role of 5alpha-androstane-3,17-dione in Polycystic Ovary Syndrome

Since 82 articles are available for 5-*α*-androstane-3,17-dione (5-*α*A), the predictions are derived from both its literature and that of its metabolic neighbourhood. The top 25 predictions ranked by *LogOdds* are presented in Supplementary Table S3, along with the p-value from a right-tailed Fisher exact test using the same data for comparison (S1.2). The highest ranked associations are both supported by several mentions of the compound and by the neighbourhood (high *priorLogOdds*). They correspond to mildly- interesting predictions as the literature of the compound alone would have been sufficient (significant Fisher p-value): the neighbourhood only strengthens the relation. Instead, we choose to focus on the relation with Polycystic Ovary Syndrome (PCOS) which has a non-significant Fisher p-value and only one article supporting the relation [32]. The *priorLogOdds* (5.47) indicates that the literature gathered from the metabolic neighbourhood seems highly related to the disease (Figure 5). While the literature of the compound alone is insufficient to highlight an association with PCOS, the posterior distribution, combining information available from the compound and its neighbours, strongly suggests one (*LogOdds* = 6.23 and *Log*_2_*FC* = 3.14). Androsterone, a direct neighbour of 5-*α*A through the reaction *3(or 17)-alpha-hydroxysteroid dehydrogenase*, is the main contributor supporting the prediction (Figure 5). Additional contributors such as testosterone, testosterone-sulfate, estradiol-17*β* and progesterone are more distant metabolically (2-3 reactions) but are also frequently mentioned in this context [33, 34, 35, 36, 37, 38, 39]. Also, PCOS is much more frequently mentioned in the literature of 4-androstene-3,17-dione compared to the other metabolites in the neighbourhood, making it an outlier among the contributors. Interestingly, its contribution significantly drops in the posterior distribution (See details in Supplementary materials S4.5 and Table S4). A view of the metabolic neighbourhood of 5-*α*A is also presented in Supplementary Figure S3.

**Figure 5:**
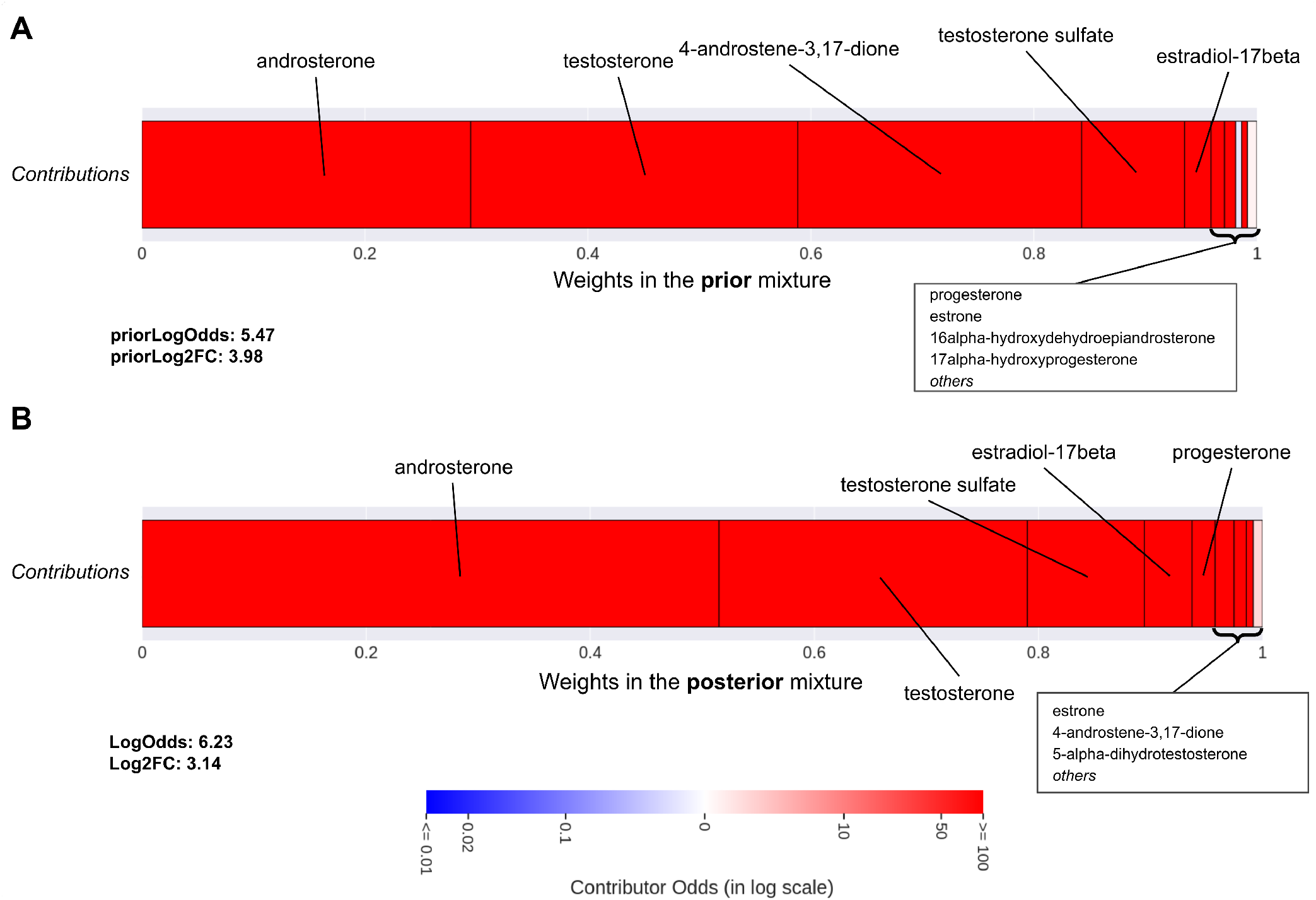
Profile of the contributors for the association between 5alpha-androstane-3,17-dione and Polycystic Ovary Syndrome in the **prior** mixture (**A**) and in the **posterior** mixture (**B**). Contributors are organised in blocks by increasing weights in the mixture from left to right, and the weights also give the width of the block. The colour of each block associated with a contributor depends on its individual *LogOdds*, from blue to red, for *negative* (less likely) to *positive* (more likely) contributions respectively. Details in Table S4.

To illustrate the influence of the observations on the posterior distribution, we re-evaluated the relation by removing the single co-occurrence between the 5-*α*A and PCOS. By suppressing this mention, the *LogOdds* drops to 3.67, *Log*_2_*FC* to 2.80, and the weights in the posterior mixture change according to the new observations (See Supplementary Figure S2). For instance the weight of androsterone, which literature mentions PCOS less frequently than the other top contributors (testosterone, estradiol, etc.), increased while those of the others decreased. More significantly, the weight of 16alphahydroxydehydroepiandrosterone, which is never mentioned with the disease, increases from 0.38% to 3%. More details in Supplementary materials S4.5. By removing this mention, the likelihood of the evidences for each contributor changed, favouring those for whom the disease is less likely to be mentioned in an article. Although the relation is still suggested by the neighbourhood, this result shows the impact of the available literature on the predictions.

## 4 Discussion

The interpretation of experimental results in metabolomics requires an intensive dive in the scientific literature. In a biomedical context, researchers often seek studies that mention metabolites from an observed signature, as well as report variations in their concentration in similar phenotypes. However, we have shown that there is a strong imbalance in the distribution of the literature among metabolites, suggesting that this research could be restricted to a subset of the initial metabolic signature. Even if this imbalance is accentuated by technical limitations, it also reflects biological facts: some metabolites are more central and sensitive to phenotypic alterations and would therefore be more frequently reported. Nonetheless, they do not necessarily provide key information when interpreting results, because they do not point to dysregulations on specific pathways. To extend the available data to help interpret results, we propose a method to suggest relations between understudied metabolites and diseases. Most metabolites (62%) in the network have no literature available, and many cannot be mapped to their corresponding PubChem identifier. It is a common issue when dealing with metabolic networks, as they are initially built for modelling purposes [40]. The absence of annotations also indicates that a compound is not widely described and studied, which may suggest that little literature has actually been lost.

The predictions for metabolites without literature are solely based on their prior distribution which is built from a mixture of the neighbouring literature. We first evaluated the prior alone on a validation dataset (AUC ≈ 0.78) and showed that it holds relevant information about the biomedical context of metabolites. Since the contributors, their weights, and influences in the mixture distribution (more or less likely to mention the disease in an article) are known, the prior is transparent by design. In the example of hydroxytyrosol, the prediction was mainly derived from the literature of dopamine, 3-4-dihydroxyphenylacetaldehyde (DOPAL), 3,4-dihydroxyphenylacetate (DOPAC) who all frequently mention Parkinson’s disease in their literature. Hydroxytyrosol and its contributors belong to the dopamine degradation pathway [41]. The literature supporting the relation with Parkinson’s disease mainly discusses the production of hydrogen peroxide during dopamine degradation to DOPAL by MAO enzymes. Since DOPAL is then inactivated into either DOPAC or Hydroxytyrosol, the literature that has been propagated by the contributors is metabolically relevant for hydroxytyrosol. Indeed, [42] shows that Hydroxytyrosol can induce a negative feedback inhibition on dopamine synthesis resulting in a decrease of the oxidation rate of dopamine. By indicating which and how neighbours contributed to the predictions, the contribution profile thus adds explainability to the predictions, which we believe is an important quality of the method. It can be quickly established if there was a clear consensus in the neighbourhood or if the association was only carried by one dominant contributor. In the case of *positive* suggestions, the associated literature of each contributor could be examined to understand the nature of their relation with the disease and assess the consistency of the prediction. Typically, we want to evaluate whether the relationship between the contributors and the disease can indeed be transferred to the target compound, whether it may suggest another, or whether it is irrelevant.

While a consensus is of course preferred (not matter the outcome of the prediction), some contributors may also have divergent literature for a particular disease. To complete the example of hydroxytyrosol, we show the profile of the contributors for the relation between 5-S-Cysteinyldopamine (CysDA) and Parkinson’s disease (See Figure S4.A). CysDA is the S-conjugate of dopamine and cysteine and its prior is mainly influenced by the literature of both of these precursors, at 51% and 45% respectively. While dopamine is strongly related to the disease, cysteine is much less mentioned in this context and the prior is consequently indecisive (*priorLogOdds ≈* 0.1). In this case, only the observed literature of CysDA can reduce the uncertainty by updating the prior distribution. In FORUM, 11 articles out of 33 mention CysDA and Parkinson’s disease, which has an important impact on the weights in the posterior mixture in favour of dopamine, which then becomes the dominant contributor (See Figure S4.B). Indeed, the posterior weights are proportional to the likelihood of the data according to the prior defined by each contributor. For CysDA, observations clearly suggest that it should be frequently mentioned with Parkinson’s disease, like dopamine, contrary to what is suggested by cysteine. The prediction is highly significant (*LogOdds* = 50.7, *Log*_2_*FC* = 3.87) as the literature of CysDA is very indicative. However, we would have already suggested the relation if only 2 articles out of 33 had mentioned the disease (see Figure S4.C). This highlights the sensibility of the method which may suggest still poorly supported relations, but which are consistent with the metabolic neighbourhood’s literature.

Likewise, the literature linking 5-*α*A to PCOS is not sufficient in quantity to statistically show a relation. From an expert’s perspective, only one qualitative article could be sufficient to justify a relation between a metabolite and a disease. But since the literature and the topics related with metabolomics are broad, highlighting these weakly supported relations could point to relevant paths of interpretation that may have been missed. The relation between 5-*α*A and PCOS is supported by only one article but is highly coherent in the metabolic neighbourhood, as androgen metabolism dysfunctions are central in this pathology [43]. As the contributors are widely studied metabolites (androsterone, testosterone, …) that also frequently mention the disease in their literature, the prior regarding the relationship is strong and strengthens the observations. We also show that after removing the only supporting article and computing the posterior distribution accordingly, the relation is still suggested but the *LogOdds* and *Log*_2_*FC* significantly drops. This illustrates the behaviour of the method, where the posterior distribution proposes a compromise between the compound’s literature and that of its contributors, giving more weight to those that are the most mentioned and for whom the observations are the most consistent. The neighbourhood literature can also help to discard suggestions that are supported by secondary or negligible mentions (See S4.6).

With FORUM’s data, relations are evaluated for both disease-specific MeSH and broader descriptors, representative of disease families such as *Neurodegenerative Diseases* (D019636). When there is no consensus among contributors at the level of specific diseases but they all belong to the same category of disorders, it could allow to suggest more coarse-grained relations. Although this increases the redundancy of the results, it makes it easier to grasp the overall biomedical context of some understudied metabolites.

## 5 Limitations

The most evident limitation of the proposed approach is that the assumption that the literature in the metabolic neighbourhood of a metabolite provides relevant prior knowledge on its biomedical context, is not always accurate. A short path of reactions can indeed have a major impact on the metabolic activity of compounds, resulting in separate biological pathways and invalidating the hypothesis. For instance, while dopamine is a derivative of tyrosine, the former is a neurotransmitter and the latter a fundamental amino acid. Their biomedical literature therefore covers very different topics, and one would not provide a good *a priori* on the other. Nonetheless, thanks to the transparency of the contributors’ profile, such irrelevant contributions can be identified and the corresponding predictions re-evaluated or discarded.

Based solely on the metabolic network, we ignore the regulatory mechanisms of biological pathways and only focus on biochemistry. We therefore assume that all paths of reactions are active and valid when propagating the literature, which is not true and may vary depending on physiological conditions. The predictions could potentially be improved by integrating a regulation layer, but this would add major complexity to the method and we choose to ignore these constraints by proposing a more general approach. Although reconstructions of the human metabolism like Human1 are constantly improving, they remain incomplete and some pathways (eg. lipids[44]) are simplified with missing or artificially created links, mainly for modelling purposes.

With their overflowing literature, overstudied metabolites (amino acids, cholesterol, etc.) can erase the contributions of other neighbours in the construction of a prior. This results in a strong prior which is only fuelled by the literature of one dominant contributor, and in the case of a metabolite without literature, predictions will therefore be solely based on it. We therefore provide diagnostic indicators like *Entropy, CtbAvgDistance* and *CtbAvgCorporaSize* (See S1.3) to identify these unbalanced priors and flag these predictions. Finally, a part of the biomedical literature of some influential compounds may not be related to their metabolic activity. For instance, ethanol is strongly related to bacterial infections, not as a metabolite but because of its antiseptic properties, which may suggest out-of-context relations by spreading its literature to neighbours. Although we kept it in our analysis for sake of exhaustively, it could be beneficial to remove its literature and that of metabolites with similar behaviours, for predictions on their close neighbours.

## 6 Conclusion

Based on the literature extracted from the FORUM KG, we showed the imbalance in the distribution of the literature related to metabolites. To overcome this bias, we proposed an approach in which we extend the *guilt by association* principle in the Bayesian framework. Basically, we use a mixture of the literature of the metabolic neighbourhood of a compound to build a prior distribution on the probability that one of its articles would mention a particular disease. The transparency of the contributor’s profile is essential and helps diagnose and explain the predictions by indicating which and how metabolic neighbours have contributed. More than 35,000 relations between metabolites and disease-related MeSH descriptors have been extracted and are available on the FORUM ftp. These relations may help interpret metabolic signatures when no or little information can be found in the literature or databases. In the upcoming release of the FORUM KG, these relations will be integrated as a peripheral graph to supplement the existing metabolite-disease associations and create new paths of hypotheses. In this analysis we restricted our predictions to diseases-related concept because the metabolic network, although suitable for propagating this type of relationship, would be less reliable for propagating functional relations for instance. The process is also network dependent, which means that using a different metabolic network (human or other organisms) could result in different suggestions. Nonetheless, the approach could be extended to other entities (genes, proteins) and relations, as long as the related literature is available and the neighbourhood of an individual can provide a meaningful prior. Finally, as the literature grows rapidly and metabolic networks become more comprehensive, we hope that this will also improve both the quantity and quality of the suggestions in the future.

## 7 Method

### 7.1 Settings

The approach is metabolite-centric, considering all the available literature for each metabolite and its co-mentions with disease-related MeSH descriptors as input data. Note that each article frequently mentions numerous metabolites and therefore the literature related to each metabolite, in terms of publications, is not exclusive to that chemical, but can be shared with others. We thus call a ‘mention’ the fact that an article mentions a metabolite.

For *M* metabolites in the metabolic network, we note *n*_*i*_ the total number of mentions of a metabolite *i* and then define 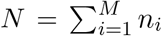 as the total number of mentions in the network. Given a specific disease-related MeSH descriptor, we also define *y*_*i*_ as the number of articles co-mentioning the metabolite *i* and the disease, with 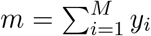 the total number of mentions involving that disease. Details on the extraction of literature data from the FORUM KG are presented in S1.2.

For a metabolite *k* of interest, the random variable *p*_*k*_ denotes the probability that an article mentioning the metabolite *k*, also mentions the disease. The aim of the method is to estimate the posterior distribution of *p*_*k*_, given a prior built from the literature of its metabolic neighbourhood. To assess the strength of their relation, *p*_*k*_ is then compared to the expected probability 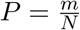 that any mentions of a metabolite in the literature also involves the disease. As in the method summary, the scenario in Figure 1 will be used to illustrate the different steps.

### 7.2 Estimating the contributions of metabolic neighbours

Based on the assumption that the literature from the metabolic neighbourhood of a compound could provide a useful prior on its biomedical context, the first step is to propagate the neighbours’ literature. A random walk with restart (RWR) algorithm (or Personalized PageRank) is used to model a mention, sent by a metabolite *i*, which moves randomly through the edges in the network and reaches another compound *k*. At each step, the mention has a probability *α*, named the *damping factor*, of continuing the walk and (1 *− α*) of restarting from the metabolite *i*. The result is a probability vector *π*_*i*._, indicating the probability that a mention sent by *i* reaches any metabolites *k* in the network, noted *π*_*i,k*_. The expected number of mentions sent by *i* that reach the compound *k* are then *π*_*i,k*_*n*_*i*_. However, in this model, a compound can receive its own mentions (*π*_*k,k*_ *>* 0) although only those derived from the neighbourhood should be used to build the prior, as the metabolite should not influence itself. A second bias is relative to the set of neighbours for which a metabolite is *allowed* to contribute to their prior. Metabolites with very large corpora (Glucose, Tryptophan, etc.) can propagate their literature to distant metabolites in the network, even if their probability to reach them is low. In the case of metabolites with a rarely mentioned direct neighbourhood, they can predominantly contribute to the prior, although they are not metabolically relevant. This bias is accentuated by the highly skewed distribution of the literature.

To contribute to the prior of *k*, we therefore require that a metabolite *i* should have a probability of reaching *k* (without considering the walks that land on itself) greater than the probability of choosing *k* randomly. The set of metabolites *k* to which *i* is allowed to contribute, namely the influence neighbourhood of *i*, noted *H*_*i*_, is therefore defined as :

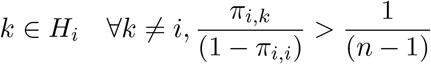

According to these probabilities, the quantity of literature sent by *i* that reaches *k* is noted *t*_*i,k*_ such as:

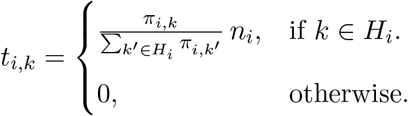

These aspects are illustrated in Figure 1.B: *B* propagates its literature to its neighbourhood but no mentions return to *B, B* is not allowed to send mentions to *Z* (being too far) and *A* receives *t*_*B,A*_ mentions from *B*. Symmetrically, we defined *T*_*k*_ as the set of contributors of *k*, such that *t*_*i,k*_ *>* 0. Each contributor *i*, has a weight *w*_*i,k*_ in the prior of *k*, representing the proportion of literature reaching *k*, that was sent by *i*:

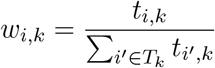

The weight vector for compound *k* is noted ***w***_***k***_. In Figure 1.C, *w*_*B,A*_ is the weight of *B* in the prior of *A* and as *A* cannot contribute to itself, *w*_*A,A*_ = 0.

### 7.3 Mixing neighbouring literature to build a prior

We assume that *a priori*, any metabolites and diseases are independent concepts in the literature, so that mention of the former does not affect the probability of mentioning the latter and *E*[*p*_*i*_] = *P*. Under this assumption, for any contributor *i*, the prior distribution of *p*_*i*_ is modelled as a Beta distribution parameterized by mean (*µ*) and sample size (*ν*):

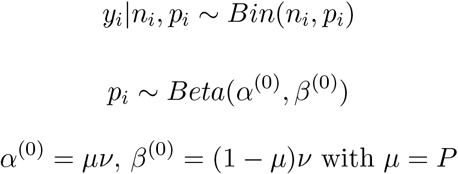

The sample size *ν* is a hyperparameter and controls the variance, the higher *ν*, the lower the variance: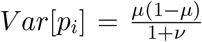. More intuitively, *ν* can be seen as the number of pseudo-obeservations that support this prior belief. The higher *ν*, the more each contributor *i* would have to bring new evidences (*n*_*i*_) to change the prior belief[45]. As the Beta distribution is a conjugate prior of the Binomial distribution, the posterior distribution of *p*_*i*_ can also be expressed as a Beta distribution:

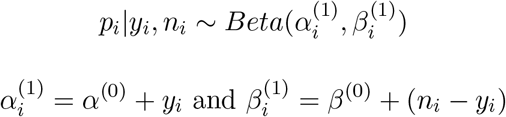

For overlooked neighbours which might bring unreliable contributions, the posterior distribution of *p*_*i*_ acts as a shrinkage procedure, by adjusting the probability distribution toward the overall probability *P* of mentioning the disease. This is illustrated in Figure 1.D: the contributor *F* has only 2 annotated publications, with one mentioning the disease. While the raw estimated probability that F mentions the disease clearly seems overestimated due to its small amount of available literature, the posterior distribution of *p*_*F*_ is more reliable.

As illustrated in Figure 1.E, the prior distribution of *p*_*k*_, also noted *f*_*prior*_, is then defined as a mixture of the distributions 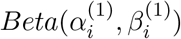 of each contributor, weighted by *w*_*i,k*_:

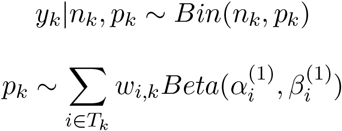

In summary, the parameters *α* and *ν* respectively control the average distance to which a metabolite is allowed to contribute to the prior of its neighbours, and the strength of the initial prior in the shrinkage procedure. The impact of these parameters on the constructed prior and predictions is discussed in S4.1. In the analyses presented in 3.3 and 3.4, we set *α* = 0.4 and *ν* = 1000.

### 7.4 Updating prior and selecting novel associations

For the compound *k*, the final posterior mixture distribution of *p*_*k*_, also noted *f*_*post*_ (Cf. Figure 1.F), is thus expressed as a mixture of the updated posterior distributions of each contributor, reweighted according to the observed data (*n*_*k*_ and *y*_*k*_):

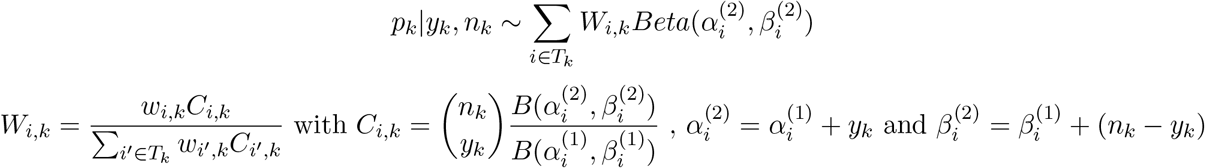

*C*_*i,k*_ represents the probability of observing the data (*y*_*k*_, *n*_*k*_) of the metabolite *k*, where *p*_*k*_ is drawn from the Beta distribution of the contributor 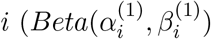, as in a Beta-binomial model. Therefore, the posterior weights in the mixture (*W*_*i,k*_) correspond to the initial weights (*w*_*i,k*_), reweighted according to the likelihood of the observations from the perspective of the contributor *i*.

From the mixture distribution, we evaluate the probability that *p*_*k*_ ≤*P*, or the posterior error that an article mentioning the metabolite *k*, would mention the disease more frequently than expected, noted *CDF*. We set *q* = 1 − *CDF* and then use the log odds of *q*, such as 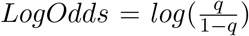Therefore, if *LogOdds* > 0, it is more likely that the metabolite *k* is related to the MeSH than it is not, and viceverca. Also, we defined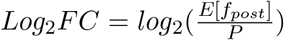 As *LogOdds* can lead to infinite values (if CDF wasn’t precisely computed and approximated to 0), the *Log*_2_*FC* can in turn provide a useful estimator to rank the relations. In turn, *Log*_2_*FC* is much more sensitive to outlier contributors than *LogOdds*. When evaluating predictions, *LogOdds* should be considered as a measure of significance and *Log*_2_*FC* as a measure of effect size. Finally, *LogOdds* and *Log*_2_*FC* can also be computed independently for each contributor *i* using their associated component in the prior 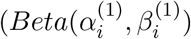 and posterior mixture 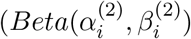.

### 7.5 Different scenarios

For metabolites mentioned in few articles and with literature available in the neighbourhood (2), the behaviour of the method is exactly as described above. When the compound *k* has no annotated articles (1), only the distribution *f*_*prior*_ is used to compute *LogOdds* and *Log*_2_*FC*. In summary, for metabolites without literature, *LogOdds* and *Log*_2_*FC* are derived from *f*_*prior*_, while for metabolites with literature, they are obtained from *f*_*post*_. For the latter, *priorLogOdds* and *priorLog*_2_*FC* are computed from the prior distribution *f*_*prior*_ and aim to represent the belief of the metabolic neighbourhood, without the influence of the compound’s literature.

There may be no literature available in the neighbourhood of some metabolites. In this case, the prior distribution is simply defined by *Beta*(*α*^(0)^, *β*^(0)^) and then the posterior distribution is 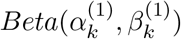. In the worst-case, where no literature is available for the metabolite and its neighbourhood, the basic distribution *Beta*(*α*^(0)^, *β*^(0)^) is used, but predictions are automatically discarded.

Since the construction of the prior from the neighbourhood’s literature is critical in the proposed method, several diagnostic values are also reported to judge its consistency. Those additional indicators are detailed in Supplementary materials S1.3.

## Acknowledgements

We are very grateful to Juliette Cooke for proofreading the manuscript.

## References

[1] Piñero, J., Ramırez-Anguita, J.M., Saüch-Pitarch, J., Ronzano, F., Centeno, E., Sanz, F., Furlong, L.I.: The DisGeNET knowledge platform for disease genomics: 2019 update. Nucleic acids research 48(D1), 845–855 (2020). Publisher: Oxford University Press

[2] UniProt Consortium, T.: UniProt: the universal protein knowledgebase. Nucleic Acids Research 46(5), 2699–2699 (2018). doi:10.1093/nar/gky092

[3] Wishart, D.S., Feunang, Y.D., Marcu, A., Guo, A.C., Liang, K., V’azquez-Fresno, R., Sajed, T., Johnson, D., Li, C., Karu, N., et al.: HMDB 4.0: the human metabolome database for 2018. Nucleic acids research 46(D1), 08–617 (2018). Publisher: Oxford University Press

[4] Mattingly, C.J., Rosenstein, M.C., Colby, G.T., Forrest Jr, J.N., Boyer, J.L.: The Comparative Toxi-cogenomics Database (CTD): a resource for comparative toxicological studies. Journal of Experimental Zoology Part A: Comparative Experimental Biology 305A(9), 689–692 (2006). doi:10.1002/jez.a.307

[5] Wishart, D.S., Bartok, B., Oler, E., Liang, K.Y.H., Budinski, Z., Berjanskii, M., Guo, A., Cao, X., Wilson, M.: MarkerDB: an online database of molecular biomarkers. Nucleic Acids Research 49(D1), 1259–1267 (2021). doi:10.1093/nar/gkaa1067

[6] Delmas, M., Filangi, O., Paulhe, N., Vinson, F., Duperier, C., Garrier, W., Saunier, P.-E., Pitarch, Y., Jourdan, F., Giacomoni, F., Frainay, C.: FORUM: building a Knowledge Graph from public databases and scientific literature to extract associations between chemicals and diseases. Bioinformatics 37(21), 3896–3904 (2021). doi:10.1093/bioinformatics/btab627

[7] Bornmann, L., Haunschild, R., Mutz, R.: Growth rates of modern science: a latent piecewise growth curve approach to model publication numbers from established and new literature databases. Humanities and Social Sciences Communications 8(1), 1–15 (2021). doi:10.1057/s41599-021-00903-w. Number: 1 Publisher: Palgrave

[8] Su, A.I., Hogenesch, J.B.: Power-law-like distributions in biomedical publications and research funding. Genome Biology 8(4), 404 (2007). doi:10.1186/gb-2007-8-4-404

[9] Edwards, A.M., Isserlin, R., Bader, G.D., Frye, S.V., Willson, T.M., Yu, F.H.: Too many roads not taken. Nature 470(7333), 163–165 (2011). doi:10.1038/470163a

[10] Wood, V., Lock, A., Harris, M.A., Rutherford, K., Bähler, J., Oliver, S.G.: Hidden in plain sight: what remains to be discovered in the eukaryotic proteome? Open Biology 9(2), 180241 (2019). doi:10.1098/rsob.180241

[11] Pandey, A.K., Lu, L., Wang, X., Homayouni, R., Williams, R.W.: Functionally Enigmatic Genes: A Case Study of the Brain Ignorome. PLoS ONE 9(2), 88889 (2014). doi:10.1371/journal.pone.0088889

[12] Stoeger, T., Gerlach, M., Morimoto, R.I., Nunes Amaral, L.A.: Large-scale investigation of the reasons why potentially important genes are ignored. PLOS Biology 16(9), 2006643 (2018). doi:10.1371/journal.pbio.2006643

[13] Perc, M.: The Matthew effect in empirical data. Journal of The Royal Society Interface 11(98), 20140378 (2014). doi:10.1098/rsif.2014.0378

[14] Miga, K.H., Koren, S., Rhie, A., Vollger, M.R., Gershman, A., Bzikadze, A., Brooks, S., Howe, E., Porubsky, D., Logsdon, G.A., Schneider, V.A., Potapova, T., Wood, J., Chow, W., Armstrong, J., Fredrickson, J., Pak, E., Tigyi, K., Kremitzki, M., Markovic, C., Maduro, V., Dutra, A., Bouffard, G.G., Chang, A.M., Hansen, N.F., Wilfert, A.B., Thibaud-Nissen, F., Schmitt, A.D., Belton, J.-M., Selvaraj, S., Dennis, M.Y., Soto, D.C., Sahasrabudhe, R., Kaya, G., Quick, J., Loman, N.J., Holmes, N., Loose, M., Surti, U., Risques, R.a., Graves Lindsay, T.A., Fulton, R., Hall, I., Paten, B., Howe, K., Timp, W., Young, A., Mullikin, J.C., Pevzner, P.A., Gerton, J.L., Sullivan, B.A., Eichler, E.E., Phillippy, A.M.: Telomere-to-telomere assembly of a complete human X chromosome. Nature 585(7823), 79–84 (2020). doi:10.1038/s41586-020-2547-7

[15] Fiehn, O.: Metabolomics — the link between genotypes and phenotypes. In: Town, C. (ed.) Functional Genomics, pp. 155–171. Springer, Dordrecht (2002). doi:10.1007/978-94-010-0448-011. http://link.springer.com/10.1007/978-94-010-0448-011

[16] Gertsman, I., Barshop, B.A.: Promises and pitfalls of untargeted metabolomics. Journal of Inherited Metabolic Disease 41(3), 355–366 (2018). doi:10.1007/s10545-017-0130-7

[17] Ivanisevic, J., Want, E.J.: From Samples to Insights into Metabolism: Uncovering Biologically Relevant Information in LC-HRMS Metabolomics Data. Metabolites 9(12), 308 (2019). doi:10.3390/metabo9120308

[18] da Silva, R.R., Dorrestein, P.C., Quinn, R.A.: Illuminating the dark matter in metabolomics. Proceedings of the National Academy of Sciences 112(41), 12549–12550 (2015). doi:10.1073/pnas.1516878112

[19] Dias, D., Jones, O., Beale, D., Boughton, B., Benheim, D., Kouremenos, K., Wolfender, J.-L., Wishart, D.: Current and Future Perspectives on the Structural Identification of Small Molecules in Biological Systems. Metabolites 6(4), 46 (2016). doi:10.3390/metabo6040046

[20] Kim, S., Thiessen, P.A., Cheng, T., Yu, B., Shoemaker, B.A., Wang, J., Bolton, E.E., Wang, Y., Bryant, S.H.: Literature information in PubChem: associations between PubChem records and scientific articles. Journal of Cheminformatics 8(1), 32 (2016). doi:10.1186/s13321-016-0142-6

[21] Lacroix, V., Cottret, L., Thebault, P., Sagot, M.-F.: An Introduction to Metabolic Networks and Their Structural Analysis. IEEE/ACM Transactions on Computational Biology and Bioinformatics 5(4), 594–617 (2008). doi:10.1109/TCBB.2008.79

[22] Robinson, J.L., Kocabaş, P., Wang, H., Cholley, P.-E., Cook, D., Nilsson, A., Anton, M., Ferreira, R., Domenzain, I., Billa, V., Limeta, A., Hedin, A., Gustafsson, J., Kerkhoven, E.J., Svensson, L.T., Palsson, B.O., Mardinoglu, A., Hansson, L., Uhl’en, M., Nielsen, J.: An atlas of human metabolism. Science Signaling 13(624), 1482 (2020). doi:10.1126/scisignal.aaz1482

[23] Hristov, B.H., Chazelle, B., Singh, M.: uKIN Combines New and Prior Information with Guided Network Propagation to Accurately Identify Disease Genes. Cell Systems 10(6), 470–4793 (2020). doi:10.1016/j.cels.2020.05.008

[24] Köhler, S., Bauer, S., Horn, D., Robinson, P.N.: Walking the Interactome for Prioritization of Candidate Disease Genes. The American Journal of Human Genetics 82(4), 949–958 (2008). doi:10.1016/j.ajhg.2008.02.013

[25] Vanunu, O., Magger, O., Ruppin, E., Shlomi, T., Sharan, R.: Associating Genes and Protein Complexes with Disease via Network Propagation. PLoS Computational Biology 6(1), 1000641 (2010). doi:10.1371/journal.pcbi.1000641

[26] Frainay, C., Aros, S., Chazalviel, M., Garcia, T., Vinson, F., Weiss, N., Colsch, B., Sedel, F., Thabut, D., Junot, C., Jourdan, F.: MetaboRank: network-based recommendation system to interpret and enrich metabolomics results. Bioinformatics 35(2), 274–283 (2019). doi:10.1093/bioinformatics/bty577

[27] Ghaderinezhad, F., Ley, C.: On the Impact of the Choice of the Prior in Bayesian Statistics. In: Tang, N. (ed.) Bayesian Inference on Complicated Data. IntechOpen, Rijeka (2020). doi:10.5772/intechopen.88994. https://www.intechopen.com/books/bayesian-inference-on-complicated-data/on-the-impact-of-the-choice-of-the-prior-in-bayesian-statistics

[28] Newman, M.E.: Power laws, Pareto distributions and Zipf’s law. Contemporary physics 46(5), 323– 351 (2005). Publisher: Taylor & Francis

[29] O’Dowd, Y., Driss, F., Dang, P.M.-C., Elbim, C., Gougerot-Pocidalo, M.-A., Pasquier, C., El-Benna, J.: Antioxidant effect of hydroxytyrosol, a polyphenol from olive oil: scavenging of hydrogen peroxide but not superoxide anion produced by human neutrophils. Biochemical Pharmacology 68(10), 2003– 2008 (2004). doi:10.1016/j.bcp.2004.06.023

[30] Monroy-Noyola, A.: Hydroxytyrosol inhibits MAO isoforms and prevents neurotoxicity inducible by MPP invivo. Frontiers in Bioscience 12(1), 25–37 (2020). doi:10.2741/s538

[31] Brunetti, G., Di Rosa, G., Scuto, M., Leri, M., Stefani, M., Schmitz-Linneweber, C., Calabrese, V., Saul, N.: Healthspan Maintenance and Prevention of Parkinson’s-like Phenotypes with Hydroxytyrosol and Oleuropein Aglycone in C. elegans. International Journal of Molecular Sciences 21(7), 2588 (2020). doi:10.3390/ijms21072588

[32] Agarwal, S.K., Judd, H.L., Magoffin, D.A.: A mechanism for the suppression of estrogen production in polycystic ovary syndrome. The Journal of Clinical Endocrinology & Metabolism 81(10), 3686–3691 (1996). doi:10.1210/jcem.81.10.8855823

[33] Xu, X.-L., Deng, S.-L., Lian, Z.-X., Yu, K.: Estrogen Receptors in Polycystic Ovary Syndrome. Cells 10(2), 459 (2021). doi:10.3390/cells10020459

[34] Matteri, R.K., Stanczyk, F.Z., Gentzschein, E.E., Delgado, C., Lobo, R.A.: Androgen sulfate and glucuronide conjugates in nonhirsute and hirsute women with polycystic ovarian syndrome. American Journal of Obstetrics and Gynecology 161(6), 1704–1709 (1989). doi:10.1016/0002-9378(89)90954-X

[35] Song, Y., Ye, W., Ye, H., Xie, T., Shen, W., Zhou, L.: Serum testosterone acts as a prognostic indicator in polycystic ovary syndrome-associated kidney injury. Physiological Reports 7(16) (2019). doi:10.14814/phy2.14219

[36] Consortium, T.E.C.A., Ruth, K.S., Day, F.R., Tyrrell, J., Thompson, D.J., Wood, A.R., Mahajan, A., Beaumont, R.N., Wittemans, L., Martin, S., Busch, A.S., Erzurumluoglu, A.M., Hollis, B., O’Mara, T.A., McCarthy, M.I., Langenberg, C., Easton, D.F., Wareham, N.J., Burgess, S., Murray, A., Ong, K.K., Frayling, T.M., Perry, J.R.B.: Using human genetics to understand the disease impacts of testosterone in men and women. Nature Medicine 26(2), 252–258 (2020). doi:10.1038/s41591-020-0751-5

[37] Doldi, N., Gessi, A., Destefani, A., Calzi, F., Ferrari, A.: Polycystic ovary syndrome: anomalies in progesterone production. Human Reproduction 13(2), 290–293 (1998). doi:10.1093/humrep/13.2.290

[38] O’Reilly, M.W., Taylor, A.E., Crabtree, N.J., Hughes, B.A., Capper, F., Crowley, R.K., Stewart, P.M., Tomlinson, J.W., Arlt, W.: Hyperandrogenemia Predicts Metabolic Phenotype in Polycystic Ovary Syndrome: The Utility of Serum Androstenedione. The Journal of Clinical Endocrinology & Metabolism 99(3), 1027–1036 (2014). doi:10.1210/jc.2013-3399

[39] Stener-Victorin, E., Holm, G., Labrie, F., Nilsson, L., Janson, P.O., Ohlsson, C.: Are there any sensitive and specific sex steroid markers for polycystic ovary syndrome? The Journal of Clinical Endocrinology and Metabolism 95(2), 810–819 (2010). doi:10.1210/jc.2009-1908

[40] Haraldsd’ottir, H.S., Thiele, I., Fleming, R.M.: Comparative evaluation of open source software for mapping between metabolite identifiers in metabolic network reconstructions: application to Recon 2. Journal of Cheminformatics 6(1), 2 (2014). doi:10.1186/1758-2946-6-2

[41] Meiser, J., Weindl, D., Hiller, K.: Complexity of dopamine metabolism. Cell communication and signaling: CCS 11(1), 34 (2013). doi:10.1186/1478-811X-11-34

[42] Goldstein, D.S., Jinsmaa, Y., Sullivan, P., Holmes, C., Kopin, I.J., Sharabi, Y.: 3,4-Dihydroxyphenylethanol (Hydroxytyrosol) Mitigates the Increase in Spontaneous Oxidation of Dopamine During Monoamine Oxidase Inhibition in PC12 Cells. Neurochemical Research 41(9), 2173– 2178 (2016). doi:10.1007/s11064-016-1959-0

[43] Nisenblat, V., Norman, R.J.: Androgens and polycystic ovary syndrome. Current Opinion in Endocrinology, Diabetes & Obesity 16(3), 224–231 (2009). doi:10.1097/MED.0b013e32832afd4d

[44] Poupin, N., Vinson, F., Moreau, A., Batut, A., Chazalviel, M., Colsch, B., Fouillen, L., Guez, S., Khoury, S., Dalloux-Chioccioli, J., et al.: Improving lipid mapping in Genome Scale Metabolic Networks using ontologies. Metabolomics 16(4), 1–11 (2020). Publisher: Springer

[45] Kruschke, J.: Doing Bayesian Data Analysis: A Tutorial with R, JAGS, and Stan. Academic Press, Boston (2014)

