## Supplementary Material for "Suggesting disease associations for overlooked metabolites using literature from metabolic neighbours"

#### S1 Supplementary methods

##### S1.1 Defining metabolic neighbourhood from metabolic network

Human1 [1] is a genome-scale reconstruction of the human metabolic network containing 13082 reactions and 8378 metabolites and is currently one of its most comprehensive representations. Even if metabolic networks are primarily designed for modelling purposes (eg. Flux analysis), they also hold valuable topological information on the human metabolism. In the aim to use this network as a support to propagate literature, items dedicated to modelling (biomass function, transport reactions, cellular compartments, etc ...) are not useful. A critical aspect is also the presence of side compounds, cofactors of reactions [2], (eg. ATP, NAD, NADP, CoA, etc ...) that could create spurious shortcuts between compounds actually very distant in the metabolic network. In addition to being hubs in the original networks, such metabolites are also widely indexed in the literature and are among the top cited metabolites. Nonetheless, their literature is scattered and they are generally not the focus of the publications that mention them, which makes them irrelevant contributors for their metabolic neighbours. To overcome this, we chose to build a carbon skeleton graph with the GSAM tool (https://forgemia.inra.fr/metexplore/gsam). We used the SBML of the metabolic network and the available SMILES annotations of metabolites for the description of their chemical structure. Monocarbon compounds (such as CO2 or formate) were also removed from the network. Based on an atom mapping procedure achieved with the RDT library [3], the rebuilt network is a compound graph connecting two compounds when they are involved in at least one reaction, each on one side, by sharing at least one carbon. Atom mapping with the RDT library was previously applied on Recon3D [4].

An illustrated example of this procedure is presented in Figure S1. In the raw metabolic network, the galactokinase reaction connects each of its substrates (Galactose, ATP) to each of its products (Galactose-1-phosphate, ADP, H+). However, while a link between Galactose and Galactose-1-phosphate or between ATP and ADP appears clearly, a direct connection between Galactose and ATP is more spurious because ATP acts as a cofactor[1] in this reaction. In this way, it seems reasonable that the Galactose-1-phosphate could receive and use as a prior knowledge the literature from Galactose, since one is structurally derived from the other, but not from ATP. Also, as ATP is involved in hundreds of reactions, this would allow many compounds, yet separated by many reactions, to share their literature with Galactose-1-phosphate. After reconstruction of the carbon skeleton graph, metabolites duplicated in several cellular compartments have also been merged in one super-compartment to provide a unified network. Only the largest component with 2704 metabolites and 10024 edges is conserved for the subsequent analysis. The carbon skeleton graph is treated as undirected to use the links substrates-products or products-substrates equivalently in the propagation process. The transition matrix $P$ was then built according to the weight policy defined in [5]. Briefly, it determines transition probabilities between compounds while accounting for the reaction level, which would otherwise be omitted by simply using the compound graph.

[1]A cofactor is a compound required to catalyse a reaction

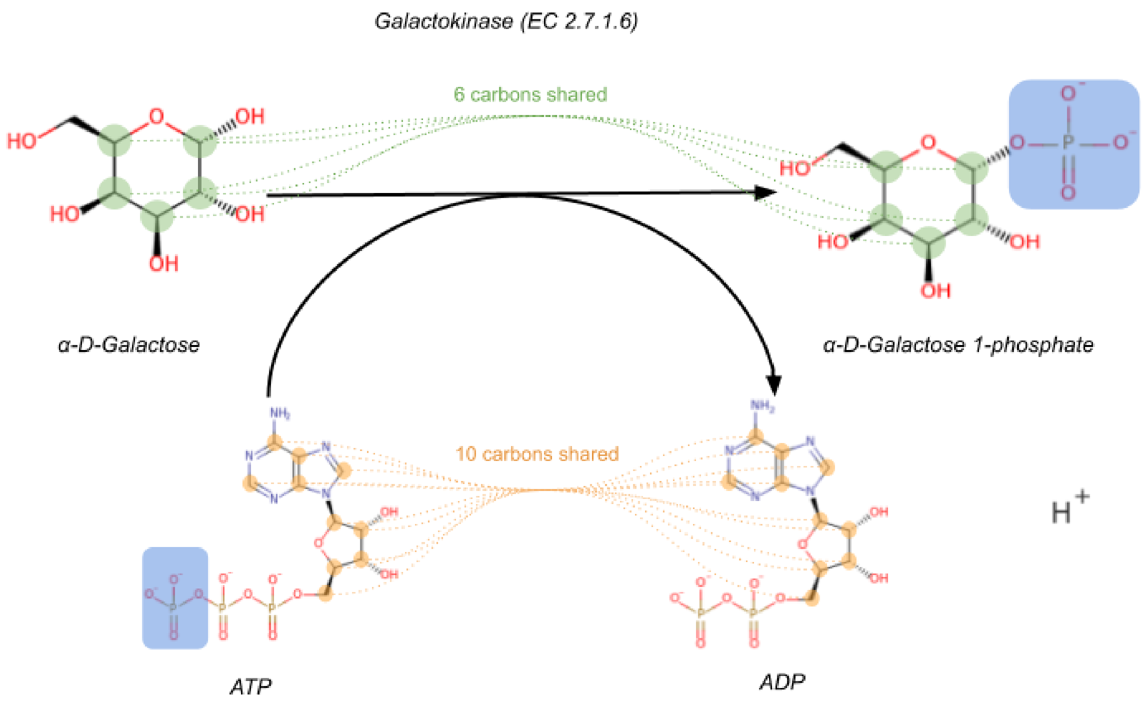


Figure S1: Example of the galactokinase reaction in the reconstruction process of the carbon skeleton graph: The galactokinase is an enzyme that catalyses the phosphorylation of galactose into galactose-1-phosphate. Coloured circles describe the carbons shared between each participant of the reaction, their number is also indicated. The blue square shows the phosphate transferred from the ATP to the galactose. There is no carbon shared between Galactose and ADP or between ATP and galactose-1-phosphate.

#### S1.2 Defining literature data for disease-metabolite associations

We used the FORUM Knowledge Graph [6] (release 2020) to extract literature data of metabolites. The FORUM KG provides links between PubChem compounds and PubMed articles, themselves indexed with a descriptive set of MeSH descriptors. The Human-GEM metabolic network v1.7 (see `https://github.com/SysBioChalmers/Human-GEM/releases/tag/v1.7.0`), converted in RDF format using an homemade java tool (`https://services.pfem.clermont.inrae.fr/gitlab/forum/sbml2rdf`), has been integrated in the FORUM KG. The total number of articles associated with each metabolite in the network, as well as their co-mentions with MeSH descriptors associated with diseases were then determined using SPARQL requests. Like in the original article, we not only considered the literature associated with MeSH descriptors of specific diseases (eg. D010300: Parkinson's disease), but also with broader descriptors representing disease families (eg. D019636: Neurodegenerative Diseases).

#### S1.3 Diagnostic values

- Entropy: The entropy value reflects both the diversity and the balance between the different contributors of the prior. Basically, the more contributors and the more uniform the weight distribution is, the higher the Entropy. For a compound $k$, the Entropy is computed on the weights in the prior mix: $H(\boldsymbol{w_k}) = \sum_{i \in T_k} \log_2(w_{i,k}) w_{i,k}$. Entropy is null when there is only one contributor, which therefore cannot be considered a "*neighbourhood*". For two contributors, the entropy is maximum and equals 1 when their contributions are equal. Obviously, one cannot require the maximum entropy as the number of contributors increases, but Entropy $> 1$ seems a reasonable threshold to apply on the predictions. For instance for 10 contributors, Entropy $> 1$ is equivalent to considering that at least 30% of the maximum entropy ($Log2(10) = 3.32$) should be reached, which would be observed if all 10 contributors contributed equally. In our network, the maximum number of contributors is 168 and by setting Entropy $> 1$, we require at least 14% of that maximal entropy.

- *CtbAvgDistance*: The average distance of the contributors, weighted by $\boldsymbol{w_k}$.

- *CtbAvgCorporaSize*: The average corpus size of the contributors, weighted by $\boldsymbol{w_k}$.

- *NbCtb*: The number of contributors in $T_k$.

- *priorLogOdds* and *priorLog*$_2$*FC*: The *LogOdds* and *Log*$_2$*FC* computed from the prior distribution $f_{prior}$. It is only provided when the targeted metabolite has literature and represents the belief of the metabolic neighbourhood. Comparing *LogOdds* and *priorLogOdds*, or *Log*$_2$*FC* and *priorLog*$_2$*FC*, allow identifying potential divergences between the literature of the compound and that of the neighbourhood. For instance, when articles mentioning a specific metabolite frequently co-mention a disease, but that disease is never mentioned in the literature of its metabolic neighbours.

#### S2 Supplementary tables

| LogOdds | Disease | MESH |
|---|---|---|
| 9.99 | Parkinson Disease | D010300 |
| 9.96 | Synucleinopathies | D000080874 |
| 9.87 | Parkinsonian Disorders | D020734 |
| 9.59 | Basal Ganglia Diseases | D001480 |
| 9.57 | Movement Disorders | D009069 |
| 8.59 | Neurodegenerative Diseases | D019636 |
| 7.31 | Brain Diseases | D001927 |
| 7.09 | Central Nervous System Diseases | D002493 |
| 6.91 | Nervous System Diseases | D009422 |
| 5.37 | Primary Dysautonomias | D054969 |

Table S1: Top 10 disease-related MeSH suggested for hydroxytyrosol, ranked by *LogOdds*

| Contributor | corpora | cooc | LogOdds | $Log_2FC$ | weights |
|---|---|---|---|---|---|
| dopamine | 98422 | 8225 | inf | 3.87 | 0.62 |
| 3.4-dihydroxyphenylacetate | 5864 | 257 | 286.04 | 2.76 | 0.33 |
| 3.4-dihydroxyphenylacetaldehyde | 111 | 38 | 50.62 | 2.80 | 0.03 |
| homovanillate | 6477 | 377 | 514.87 | 3.18 | 0.01 |
| others | 45 | 4 | 2.35 | 0.74 | 0.01 |

Table S2: The table describes different properties of the contributors for the association between hydroxytyrosol and Parkinson's Disease: *corpora* corresponds to the total number of mentions associated with the compound; *cooc* is the number of co-occurring mentions with the disease; *LogOdds* indicates the individual *LogOdds* of the contributors in the prior mixture, same for $Log_2FC$; *weights* indicates the weight of each contributor in the prior mixture. The values in *others* corresponds to the median for the remaining contributors.

| priorLogOdds | LogOdds | Disease | cooc | p-value Fisher |
|---|---|---|---|---|
| 4.16 | 23.47 | Prostatic Diseases | 9 | 1.13e-10 |
| 5.79 | 21.19 | Genital Neoplasms, Male | 8 | 2.85e-9 |
| 4.17 | 19.82 | Prostatic Neoplasms | 8 | 1.08e-9 |
| 5.68 | 19.15 | Genital Diseases, Male | 9 | 3.91e-8 |
| 5.67 | 15.83 | Urogenital Neoplasms | 8 | 4.72e-6 |
| 5.53 | 10.56 | Hair Diseases | 2 | 2.57e-3 |
| 4.00 | 9.63 | Male Urogenital Diseases | 10 | 1.36e-3 |
| 4.47 | 9.36 | Prostatic Hyperplasia | 2 | 1.27e-3 |
| 5.66 | 8.92 | Neoplasms by Site | 10 | 1.17e-2 |
| 5.11 | 8.68 | Disorder of Sex Development, 46,XY | 1 | 1.90e-2 |
| 4.85 | 8.65 | Androgen-Insensitivity Syndrome | 1 | 1.25e-2 |
| 5.49 | 8.03 | Hirsutism | 1 | 3.42e-2 |
| 6.18 | 7.82 | Virilism | 1 | 4.80e-2 |
| 5.73 | 7.60 | Skin Diseases | 5 | 4.78e-2 |
| 5.22 | 7.07 | Breast Diseases | 3 | 4.81e-2 |
| 5.68 | 7.04 | Neoplasms | 11 | 6.85e-2 |
| 4.51 | 6.99 | Alopecia | 1 | 2.87e-2 |
| 5.71 | 6.97 | Skin and Connective Tissue Diseases | 5 | 8.70e-2 |
| 4.48 | 6.93 | Hypotrichosis | 1 | 2.91e-2 |
| 4.87 | 6.72 | Breast Neoplasms | 3 | 4.24e-2 |
| 5.47 | 6.23 | Polycystic Ovary Syndrome | 1 | 1.17e-1 |
| 5.48 | 6.06 | Ovarian Cysts | 1 | 1.30e-1 |
| 5.48 | 5.86 | Cysts | 1 | 1.64e-1 |
| 5.52 | 5.62 | Disorders of Sex Development | 1 | 1.31e-1 |
| 5.54 | 5.17 | Urogenital Abnormalities | 1 | 1.59e-1 |

Table S3: Top 25 disease-related MeSH predicted for 5-$\alpha$A, ranked by *LogOdds*. The *cooc* column indicates the number of co-occurring mentions with the disease. *p-value Fisher* refers to the p-value obtained with an over-representation analysis (Fisher right-tailed exact test) using the same literature data as used for the predictions (see S1.2)

| Contributor | corpora | cooc | prior weights | posterior weights | LogOdds | $Log_2FC$ |
|---|---|---|---|---|---|---|
| androsterone | 2348 | 45 | 0.29 | 0.51 | 49.44 | 2.59 |
| testosterone | 79421 | 2521 | 0.29 | 0.28 | inf | 3.75 |
| testosterone sulfate | 69365 | 1939 | 0.09 | 0.10 | inf | 3.56 |
| estradiol-17beta | 93909 | 1102 | 0.02 | 0.04 | inf | 2.32 |
| progesterone | 75499 | 661 | 0.01 | 0.02 | 391.09 | 1.89 |
| estrone | 11455 | 114 | 0.01 | 0.02 | 78.05 | 2.00 |
| 4-androstene-3,17-dione | 8435 | 737 | 0.25 | 0.01 | inf | 5.06 |
| 5-alpha-dihydrotestosterone | 10389 | 124 | 4.00e-3 | 0.01 | 101.38 | 2.25 |
| others | 203 | 1 | 0.01 | 0.01 | 0.71 | 0.47 |

Table S4: The table describes different properties of the contributors for the association between 5-$\alpha$A and PCOS: *corpora* corresponds to the total number of mentions associated with the compound; *cooc* is the number of co-occurring mentions with the disease; *prior weights* indicates the weight of each contributor in the prior mixture; *posterior weights* indicates the weight of each contributor in the posterior mixture; *LogOdds* indicates the individual *LogOdds* of the contributors in the posterior mixture, same for $Log_2FC$; The values in *others* correspond to the median for the remaining contributors. As in Figure5, contributors are ordered by *posterior weights*.

| Contributor | corpora | cooc | prior weights | posterior weights | LogOdds | $Log_2FC$ |
|---|---|---|---|---|---|---|
| androsterone | 2348 | 45 | 0.29 | 0.61 | 47.63 | 2.56 |
| testosterone | 79421 | 2521 | 0.29 | 0.14 | inf | 3.74 |
| testosterone sulfate | 69365 | 1939 | 0.09 | 0.06 | inf | 3.56 |
| estradiol-17beta | 93909 | 1102 | 0.02 | 0.06 | inf | 2.31 |
| progesterone | 75499 | 661 | 0.01 | 0.04 | 389.81 | 1.89 |
| estrone | 11455 | 114 | 0.01 | 0.03 | 76.66 | 1.99 |
| 16alpha-hydroxydehydroepiandrosterone | 35 | 0 | 5.00e-3 | 0.03 | -0.63 | -0.16 |
| 5-alpha-dihydrotestosterone | 10389 | 124 | 3.00e-3 | 0.01 | 99.82 | 2.24 |
| androsterone sulfate | 25 | 1 | 1.00e-3 | 0.01 | 0.42 | 0.37 |
| others | 264 | 1 | 1.00e-4 | 0.01 | 0.04 | 0.17 |

Table S5: The table describes different properties of the contributors for the association between 5-$\alpha$A and PCOS **without** the single co-mention: *corpora* corresponds to the total number of mentions associated with the compound; *cooc* is the number of co-occurring mentions with the disease; *LogOdds* indicates the individual *LogOdds* of the contributors in the posterior mixture, same for $Log_2FC$; *weights* indicates the weight of each contributor in the posterior mixture. The values in *others* correspond to the median for the remaining contributors. As in Figure5, contributors are ordered by *posterior weights*.

#### S3 Supplementary figures

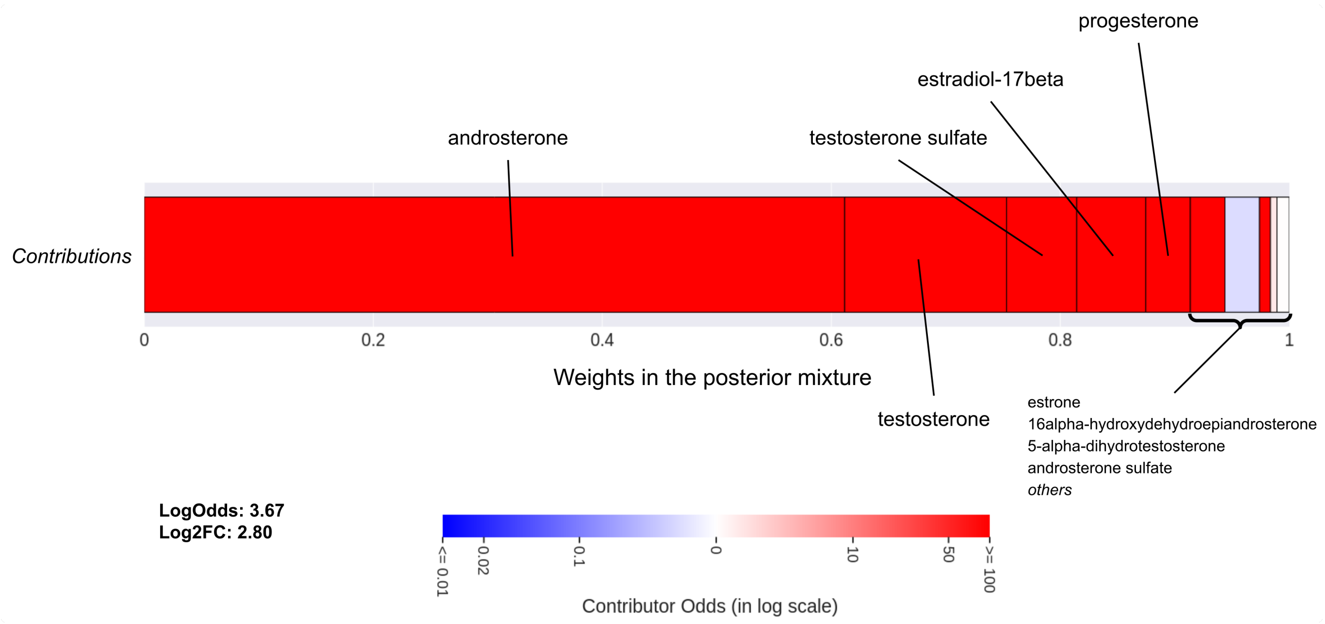


Figure S2: Profile of the contributors for the association between 5-$\alpha$A and PCOS **without** the single co-occurrence (PMID 8855823). Contributors are organised in blocks from left to right by increasing contributions. The contributions correspond to the weight of each contributor in the posterior mixture ($W_{i,k}$) and gives the width of the block. The colour of each block associated with a contributor depends on its individual *LogOdds*, from blue to red, for *negative* (less likely) to *positive* (more likely) contributions respectively. Weights and *LogOdds* are also detailed in table S5

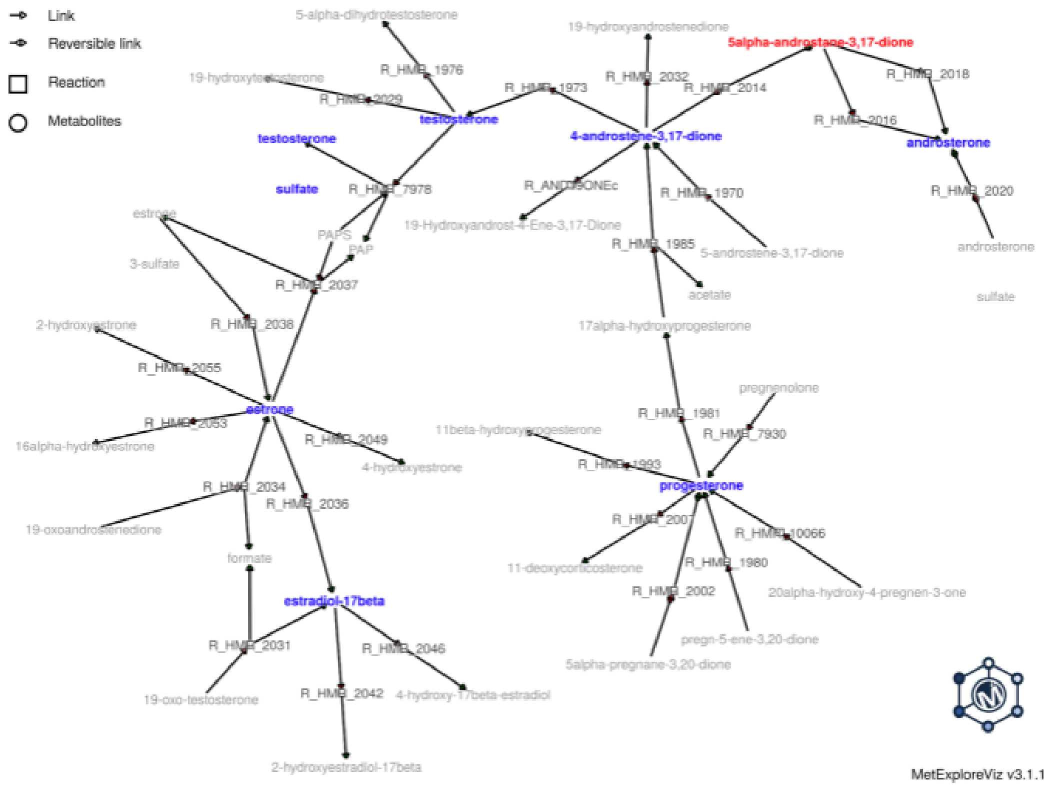


Figure S3: View of the metabolic neighbourhood of 5-$\alpha$A (in red). Main contributors of the relation with PCOS are highlighted in blue.

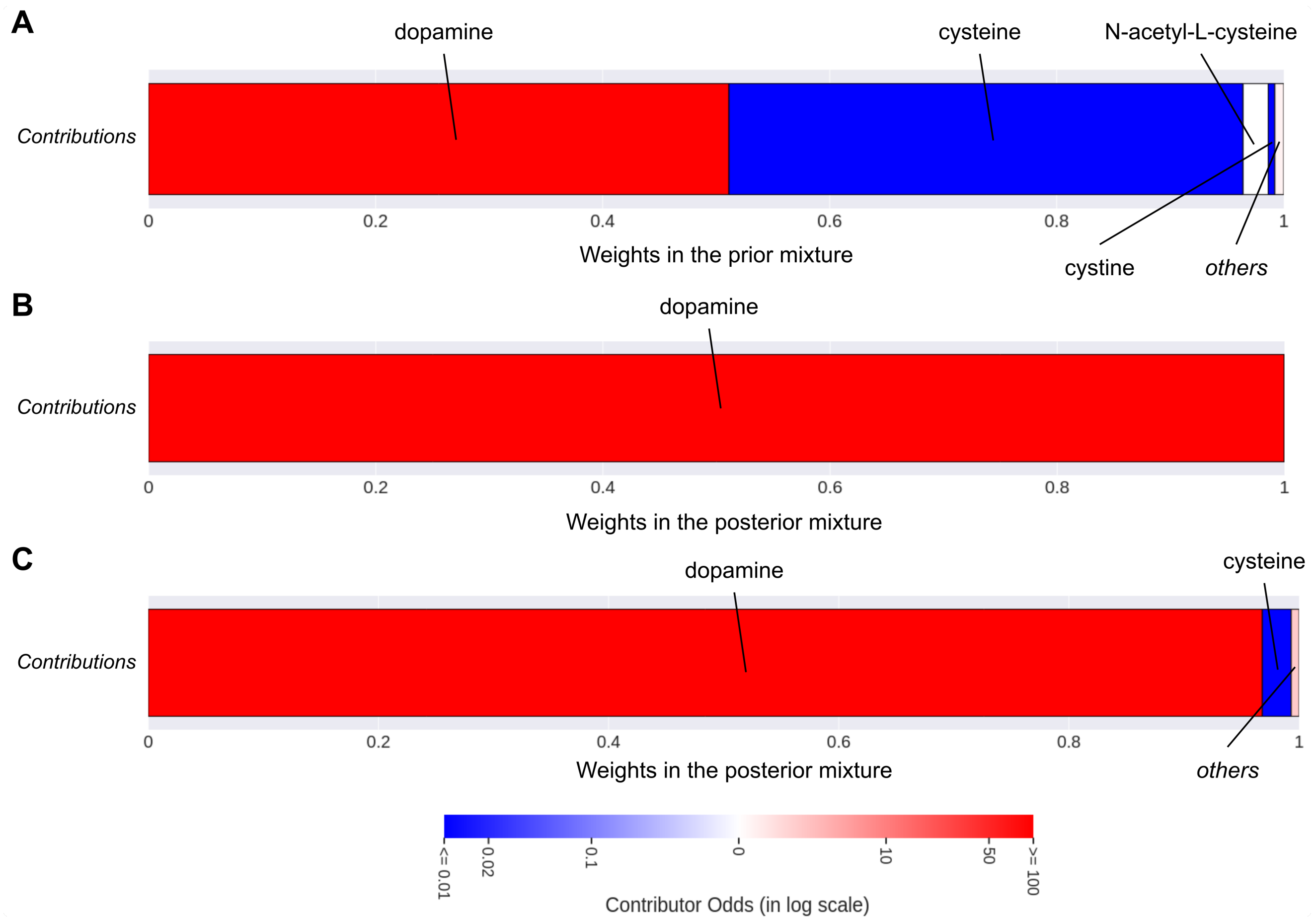


Figure S4: Profile of the contributors for the association between 5-S-Cysteinyldopamine and Parkison's disease. The profile of the contributors from the prior distribution is shown in **A** and from the posterior distribution in **B**, with actual literature data: 11 supporting articles out of 33. **C** is the profile of the contributors with only 2 co-occurrences.

### S4 Supplementary materials

#### S4.1 Damping factor $\alpha$ and theoretical sample size $\nu$: benchmark

We evaluated the impact of both hyperparameters $\alpha$ and $\nu$ on the construction of the prior and the predictions. The damping factor $\alpha$ set the probability that at each step the walk continues, so that the mention returns to its starting point with a probability $(1-\alpha)$. Increasing $\alpha$ therefore increases the average length of the walks and the radius in which a compound can propagate its literature. As the probability $\pi_{i,k}$ to reach more distant neighbours increases with $\alpha$, so do the weights $w_{i,k}$ in the prior and the average distance of the contributors (Supplementary Figure S5). At $\alpha = 0$, only the direct neighbours in the network can contribute to the prior of a compound. Considering that the direct neighbourhood is not always the richer and that we should also keep the majority of the contributors in a radius of 2 reactions, setting $\alpha$ approximately $0 < \alpha < 0.7$ seems reasonable according to the distance criteria.

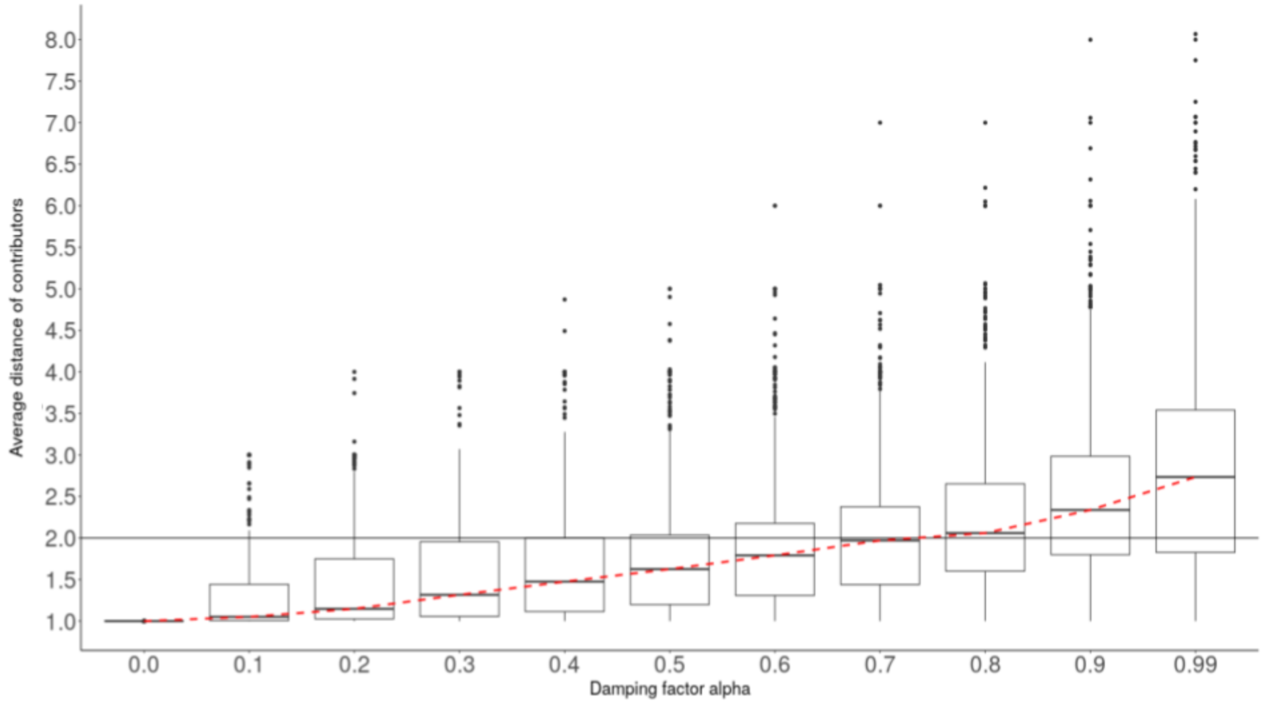


Figure S5: Boxplot of the average distance of the contributors, weighted by $w_{i,k}$, using different damping factors $\alpha$. The red dotted line connects the median of each boxplot and the black horizontal line is a threshold at an average distance of 2 reactions.

We also evaluated how the weights of the contributors distribute according to their distance to the compound, by increasing $\alpha$ (Supplementary Figure S6). As $\alpha$ increases, the closest contributors lose weight in the prior to the benefit of more distant contributors. When $\alpha > 0.7$, it seems that the contributors at a distance of at least 2 reactions became dominant in the prior. Nonetheless, it should be noted that a compound always sends more of its articles to its closest neighbours, but as this quantity decreases and the number of new contributors increases with $\alpha$, the closest neighbourhood becomes less and less influential. For $\alpha = 0.4$, we observed a relatively balanced distribution of the weights, with on average 52% of the prior derived from the direct neighbourhood, 35% from the 2-reactions neighbours, and the remaining from more distant ones.

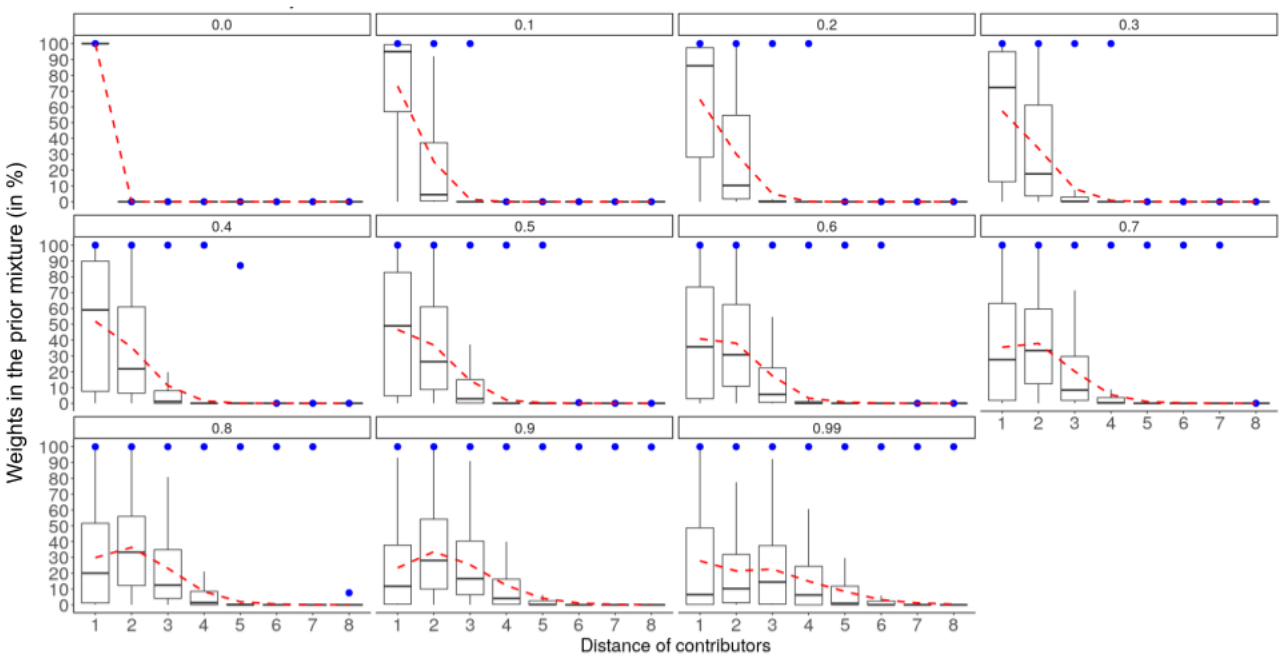


Figure S6: Distribution of the contributors' weights in the prior mixtures $w_{i,k}$, at a distance of n reactions, for several damping factors $\alpha$. The red dotted line connects the medians and the blue dots represent the maximal outliers.

Finally, we also evaluated $\alpha$ and $\nu$ using our validation dataset (S4.2) to determine their mutual impact on the built prior (Supplementary Figure S7). We have chosen to focus on the predictions based on the $LogOdds$, as the effect of the damping factor $\alpha$ is more pronounced than on the $Log_2FC$. The same methodology as described in section 3.2 was applied: only the prior distribution ($f_{prior}$) was used to compute the $LogOdds$. The TPR, FPR and precisions were computed for each combination of $\alpha$ and $\nu$ using a threshold at $LogOdds > 2$. For a fixed $\nu$, while the TPR and FPR decrease with $\alpha$, the precision increases. This suggests that as we use a closer neighbourhood, we grasp more true associations that could only be provided by the direct neighbours, but also false-positives, that would be contradicted by considering a larger neighbourhood, as there would be no consensus. For $\nu$, there is no significant impact on the predictions until $\nu = 10000$, where it shows a similar effect to that of $\alpha$ on TPR, FPR and precision. As we strengthen the initial prior with an increasing $\nu$, it will require more observations from the contributors to make it deviates from its theoretical mean set at $P$ (the overall frequency of the disease). As the median corpus size of the metabolites in the network is 172, setting a $\nu > 10000$ could erase the contributions of the majority of the compounds, unless they are highly related to the disease (high co-mention frequency). Also, setting $\nu$ to extreme values would smooth the $LogOdds$ and especially the $Log_2FC$ around 0, as the initial prior centred on $P$ would be too strong, leading to weak predictions.

However, the metabolic neighbourhood of each compound is different: some have no direct neighbours with available literature, making it necessary to use a larger neighbourhood, and some others the opposite. Therefore, there is no optimal parameters for the whole network and we can only recommend setting $0 \leq \alpha \leq 0.7$ and $1 \leq \nu \leq 10000$, increasing $\alpha$ and $\nu$ for specificity and decreasing for sensibility. From these results we therefore chose $\alpha = 0.4$ and $\nu = 1000$ in the presented analyses.

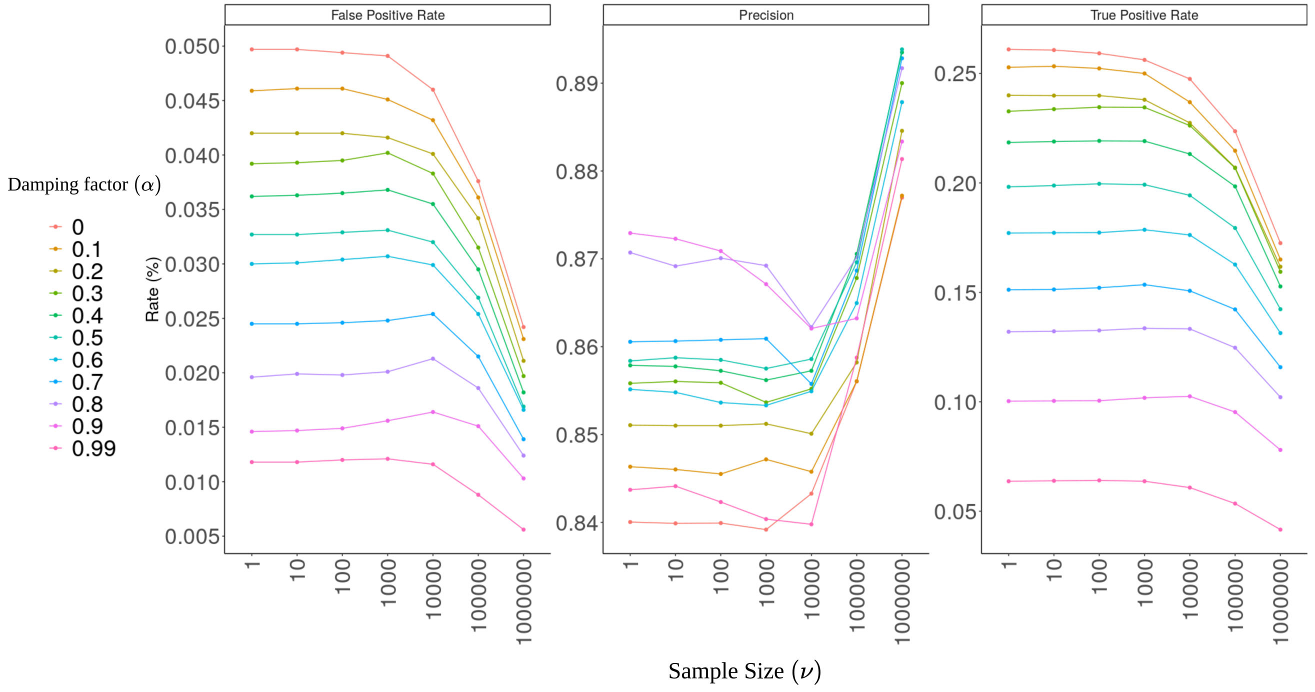


Figure S7: Evaluation of the True Positive Rate (TPR), False Positive Rate (FPR) and Precision on the validation dataset obtained with a threshold on $LogOdds > 2$, using different combinations of hyperparameters $\alpha$ and $\nu$

#### S4.2 Validation dataset

The Human-GEM metabolic network v1.7 has been converted in RDF and integrated in the FORUM KG. As in the original article, we conducted an over-representation analysis using a right-tailed fisher exact test, and extracted significant relations between metabolites and MeSH descriptors based on their co-mentions in the literature. Then, for the subset of metabolites conserved in the carbon skeleton graph (see S1.1), we randomly selected 10,000 significant relations (q-value $\leq 1e-6$ and no *weakness* [6]) with disease-related MeSH. For negative examples, we randomly generated 10,000 metabolite-MeSH pairs using the same set of metabolites and MeSH as for positives examples, ensuring that they are not positive examples if they exist (q-value $> 1e-6$). Among the 1025 metabolites with available literature in the carbon skeleton graph, 455 are present in the validation dataset.

The method has been implemented with a *forget* option, to not update the prior mixture $f_{prior}$ with the metabolite literature. *LogOdds* and $Log_2FC$ are computed directly from the prior distribution, as if the metabolite has no literature. The analysis was run on all the examples with the *forget* option and AUC and ROC curves were computed using the R library *pROC*.

##### S4.2.1 Baselines

We note $P_j = \frac{m_j}{N}$ the probability to mention the disease $j$, with $m_j$ the number of mentions involving $j$ and $N$ the total number of mentions in the metabolic network. Also, $p_{i,j} = \frac{y_{i,j}}{n_i}$ is the probability that an article mentioning the metabolite $i$, mentions the disease $j$. $y_{i,j}$ is the number of co-mentions between $i$ and $j$ and $n_i$ is the total number of articles mentioning $i$.

In Baseline-Freq, for an association between a metabolite $i$ and a disease-related MeSH $j$, the predictor is simply $P_j$, the overall probability to mention the disease.

In Baseline-DN, the predictor is the ratio between the average probability to mention the disease in the direct neighbourhood of the metabolite $i$ (noted $DN_i$) and the overall probability:

$$\frac{\sum_{u \in DN_i} \frac{p_{u,j}}{|DN_i|}}{P_j}$$

Baseline-DN is thus comparable in form to $Log_2FC$.

#### S4.3 Evaluation using simulated overlooked metabolites

To evaluate the performance of the predictions based on the posterior distribution ($f_{post}$), we build a second validation dataset with simulated overlooked metabolites. Similarly to S4.2, we extracted 10,000 significant relations with disease-related MeSH and generated 10,000 random pairs, but only for metabolites with more than 100 articles. To simulate overlooked metabolites, we generated random samples from a binomial distribution, considering the observed co-mention frequency for positive examples and the marginal frequency of mentioning the disease for negative examples (independence hypothesis). We used three different sample sizes to represent different degrees of overlooked metabolites (10, 50 and 100) and generated 10 replicates by sample size. We set $\alpha = 0.4$ and $\nu = 1000$ for the method and compared it against a new baseline (Baseline-DN+Cpd), similar to Baseline-DN, but in which the average probability includes the metabolite's literature. In contrast to Baseline-DN+Cpd, the targeted metabolite is not considered as a new contributor in the proposed approach, but is used to refine the prior distribution. The average ROC curves per tested sample sizes obtained for the method and Baseline-DN+Cpd on this new validation dataset are shown in Figure S8.A, along with the associated AUC values in Table S6. Despite its simplicity, Baseline-DN+Cpd performs well, but the proposed approach shows better performances on all the tested sample sizes.

Focusing on overlooked metabolites, the most challenging scenarios are those where positive examples apparently show no co-mentions, and conversely, when co-mentions (e.g. anecdotal) wrongly support negative examples. These configurations are common in the literature and could lead to many false negatives and false positives. We refer to them as *Hard cases*. To build validation datasets for *Hard cases*, we extracted from the previous generated simulations: all available positive examples where there is no co-occurrence, and, all the negative examples where there is at least one co-occurrence. On these *Hard cases*, the method outperforms the baseline, which, particularly on low sample sizes ($n = 10$), is misled by the observations (Figure S8 and Table S6). Despite an average AUC of roughly 0.65, the approach performs particularly well for tolerable FPR, with a TPR close to 0.3 for an FPR of 0.05 in all the tested sample sizes (Table S7). However, we can note that as the sample size increases, the examples seem to become even more complex and the performances seem to decrease slightly. Finally, these results show the robustness of the predictions for overlooked metabolites, even when the observations are misleading.

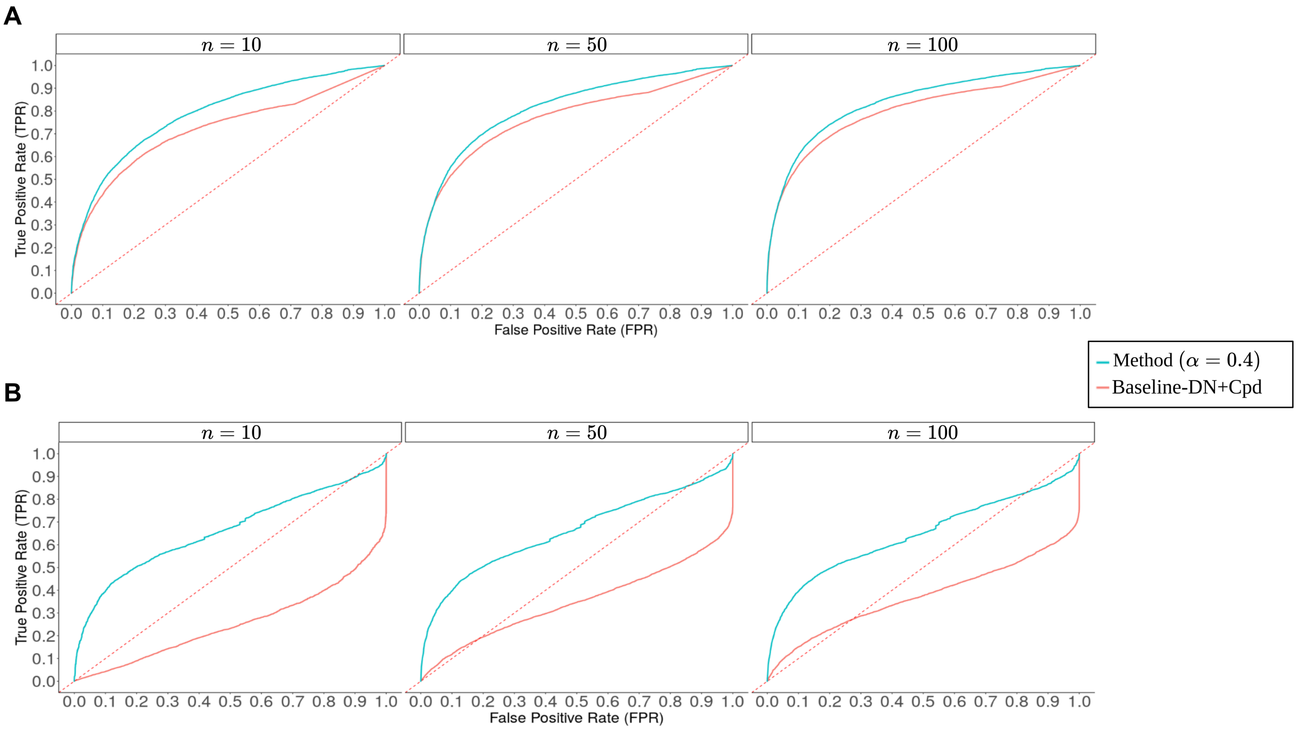


Figure S8: Average receiver operating characteristic (ROC) curves per tested sample sizes for the method set with $\alpha = 0.4$, $\nu = 1000$ and Baseline-DN+Cpd. In **A**, performances are evaluated on datasets of simulated overlooked metabolites, with increasing sample size: 10, 50 and 100. In **B**, only the *Hard cases* have been retained to evaluate the performances of the method against Baseline-DN+Cpd.

| | **Full** | | **Hard cases** | |
|---|---|---|---|---|
| Sample Size | Baseline | Method | Baseline | Method |
| n=10 | 0.72 | 0.79 | 0.25 | 0.66 |
| n=50 | 0.77 | 0.82 | 0.35 | 0.66 |
| n=100 | 0.8 | 0.84 | 0.37 | 0.64 |

Table S6: Average AUC obtained on the predictions with the proposed method and Baseline-DN+Cpd, by increasing sample sizes, on the full validation datasets (*Full*) and only on the *Hard cases*.

| **Sample Size** | **Baseline** | **Method** |
|---|---|---|
| n=10 | 0.02 | 0.29 |
| n=50 | 0.07 | 0.31 |
| n=100 | 0.1 | 0.3 |

Table S7: Average TPR on the predictions obtained with the proposed method and Baseline-DN+Cpd on *Hard cases* for an FPR fixed at 0.05 and by increasing sample sizes.

#### S4.4 Impact of the carbon skeleton graph on the predictions

Using the same validation dataset as in 3.2, we repeated the analysis on the original metabolic network (without removing spurious connections) and compared it with the carbon skeleton graph. The comparison of the ROC curves obtained from the carbon skeleton graph (CSG network) and the original Human1 metabolic network is presented in Figure S9. The AUC obtained using the original network is 0.74, significantly lower than with the CSG network (0.78). To get more insights on the impact of the CSG network on the built priors, we examined the contributions of a well-known cofactor: ATP with 109,321 annotated articles. To exclude non-significant contributions, we only consider cases where more than 10% of a metabolite's prior is represented by the literature of ATP. In the original network, ATP contributed significantly to the prior of 268 metabolites, or 6.6% of the metabolites in the network. In the CSG network, its contributions were more restricted due to the pruning of its connections as cofactor, and it only contributed significantly to the prior of 21 metabolites. These results support the use of the CSG network in the proposed approach. More generally, they illustrate the potential biases induced by cofactors when working with the topology of metabolic networks, but also the potential of the atom-mapping procedure to avoid them.

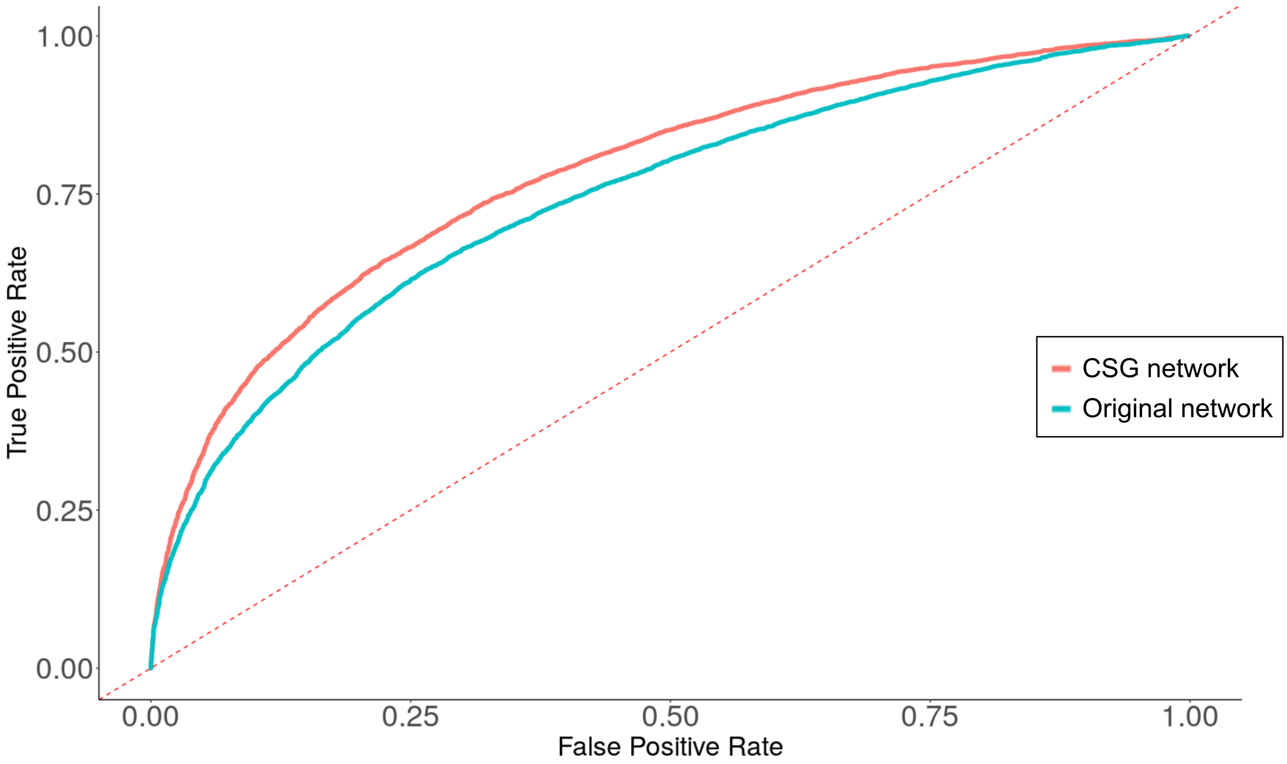


Figure S9: Receiver operating characteristic (ROC) curves using $Log_2FC$ as predictor with the carbon skeleton graph (CSG network) or the original Human1 metabolic network. The AUC are respectively 0.78 and 0.74. The red dotted line corresponds to random strategies.

##### S4.5 Comparison of the posterior distributions for 5-$\alpha$A and PCOS

In this section, we provide a more detailed view of the differences between the posterior distributions obtained with and without the co-mention. The prior mixture used is the same in both cases and a detailed view of the individual distribution of the contributors is provided in Figure S10. With an average probability of $\approx 0.08$, the 4-androstene-3,17-dione is the contributor whose literature most frequently mention the disease and therefore, the prior that it suggests is high. However, the likelihood of our observations is very low for the prior suggested by this contributor, as we would expect around 6 $(82 \times 0.08)$ co-mentions if an article mentioning 5-$\alpha$A had roughly the same probability to mention the disease. Its weight drops in the posterior distribution in both cases, as our observations are more likely compared to the literature of other contributors (Figure S11). By removing the co-mention (green on Figure S11), the weights of the contributors whose literature mentions the disease most frequently decrease, in favour of those for whom their literature mentions it less. This is illustrated by a shift to the left of the probability distribution between the red and green curves.

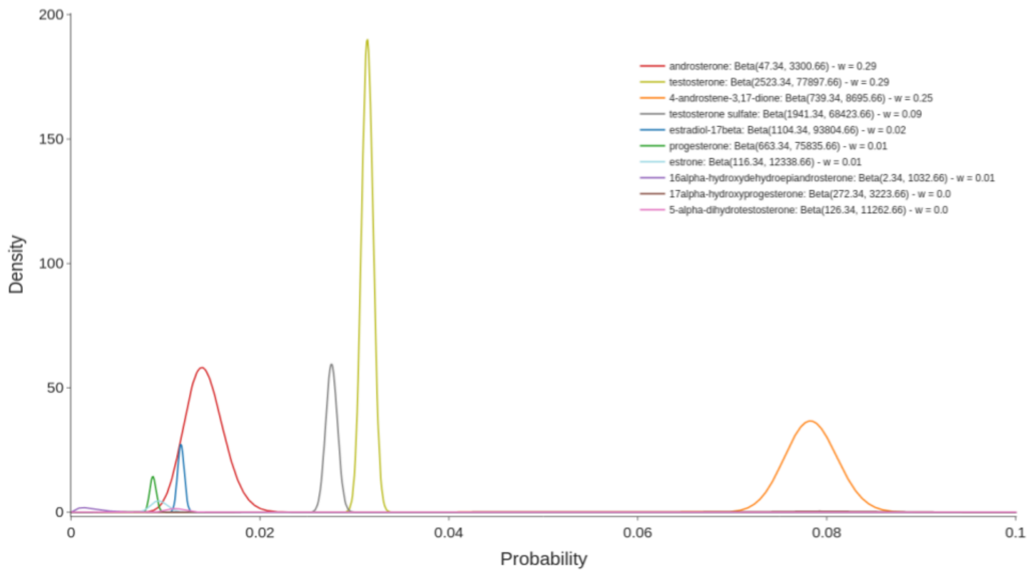


Figure S10: Detailed prior mixture of the top 10 contributors for the relation between 5-$\alpha$A and PCOS. The individual distribution of each contributor in the mixture, along the parameters of the associated *Beta* distribution, are indicated.

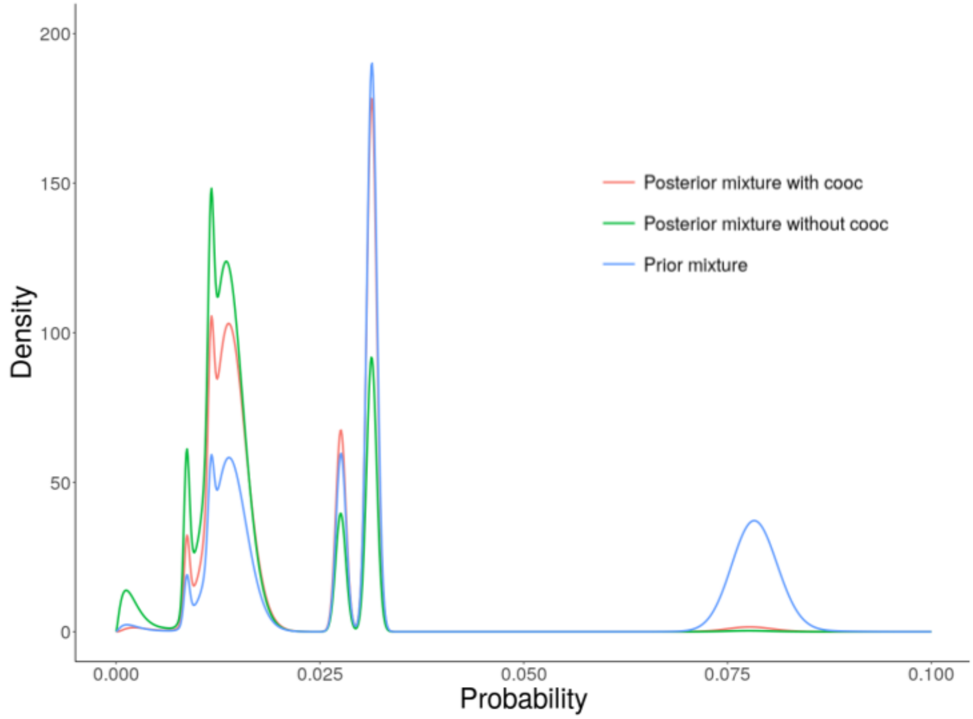


Figure S11: Pior (blue) and posterior distributions obtained with (red) and without (green) the co-mention for the relation between 5-$\alpha$A and PCOS.

##### S4.6 5-$\alpha$A and Meningioma

Like for PCOS, 5-$\alpha$A is mentioned with Meningioma in only one article but here, the fisher test would have suggested the relation (to a certain threshold: p.value $\approx$ 0.02) whereas the *LogOdds* obtained with the method is low: 0.67. Meningioma is less frequently mentioned than PCOS in the literature (824 against 10,131 annotated articles) which explains why the p.value is more significant. The Figure S12 shows the prior and posterior profiles of the contributors for the association between 5-$\alpha$A and meningioma. The literature of most of the neighbours, with the exception of progesterone, does not frequently mention the disease and therefore, the prior alone does not suggest a relation (*priorLogOdds* = -1.1). The posterior distribution logically tends to favour contributors that support the observed co-occurrence, but the resulting *LogOdds* is still low, as the observations are not sufficient to shift the prior belief. If we look at the co-mention, we see that the relation between 5-$\alpha$A and melangiomia is at least secondary in this article, as the indexing of the MeSH comes from the use of meningioma tissue as a control tissue in the conducted experiment[7]. This also shows the importance of the neighbouring literature in avoiding suggestions that could be solely derived from anecdotal mentions of rarely mentioned diseases in the literature. Of course these are suggestions, not definitive predictions, and the method returns the odds of a potential relation. Although the posterior odds don't suggest a relation, the profile shows that the progesterone, a close neighbour, seems related to the disease and therefore the association may still be worth exploring further.

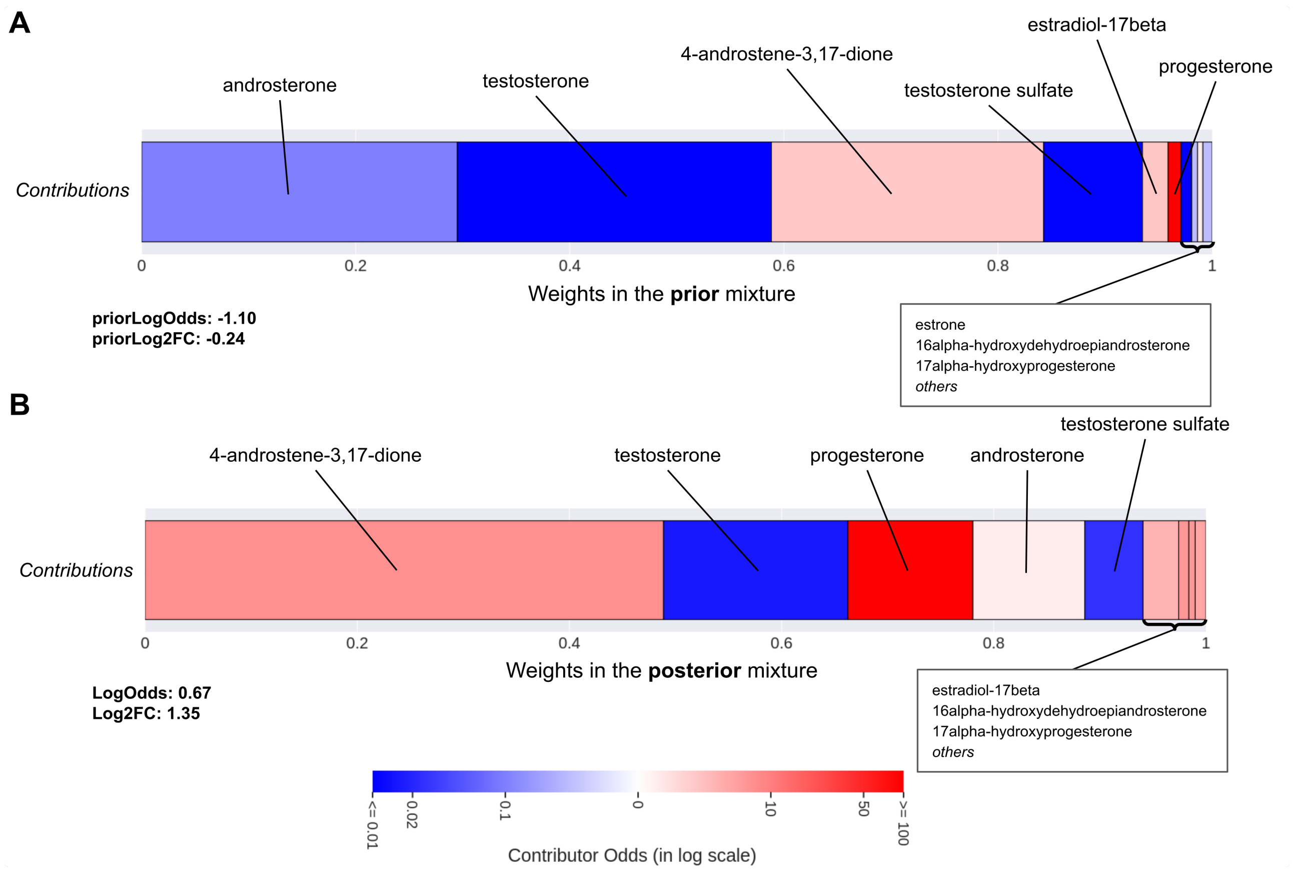


Figure S12: Profile of the contributors for the association between 5-$\alpha$A and Meningioma in the **prior** mixture (**A**) and in the **posterior** mixture (**B**). Contributors are organised in blocks by increasing weights in the mixture from left to right, and the weights also give the width of the block. The colour of each block associated with a contributor depends on its individual *LogOdds*, from blue to red, for *negative* (less likely) to *positive* (more likely) contributions respectively.

#### Supplementary References

[1] Robinson, J.L., Kocabaş, P., Wang, H., Cholley, P.-E., Cook, D., Nilsson, A., Anton, M., Ferreira, R., Domenzain, I., Billa, V., Limeta, A., Hedin, A., Gustafsson, J., Kerkhoven, E.J., Svensson, L.T., Palsson, B.O., Mardinoglu, A., Hansson, L., Uhlén, M., Nielsen, J.: An atlas of human metabolism. Science Signaling **13**(624), 1482 (2020). doi:10.1126/scisignal.aaz1482

[2] Frainay, C., Jourdan, F.: Computational methods to identify metabolic sub-networks based on metabolomic profiles. Briefings in Bioinformatics **18**(1), 43–56 (2017). doi:10.1093/bib/bbv115

[3] Rahman, S.A., Torrance, G., Baldacci, L., Martínez Cuesta, S., Fenninger, F., Gopal, N., Choudhary, S., May, J.W., Holliday, G.L., Steinbeck, C., Thornton, J.M.: Reaction Decoder Tool

(RDT): extracting features from chemical reactions. Bioinformatics **32**(13), 2065–2066 (2016). doi:10.1093/bioinformatics/btw096

[4] Preciat Gonzalez, G.A., El Assal, L.R.P., Noronha, A., Thiele, I., Haraldsdóttir, H.S., Fleming, R.M.T.: Comparative evaluation of atom mapping algorithms for balanced metabolic reactions: application to Recon 3D. Journal of Cheminformatics **9**(1), 39 (2017). doi:10.1186/s13321-017-0223-1

[5] Frainay, C., Aros, S., Chazalviel, M., Garcia, T., Vinson, F., Weiss, N., Colsch, B., Sedel, F., Thabut, D., Junot, C., Jourdan, F.: MetaboRank: network-based recommendation system to interpret and enrich metabolomics results. Bioinformatics **35**(2), 274–283 (2019). doi:10.1093/bioinformatics/bty577

[6] Delmas, M., Filangi, O., Paulhe, N., Vinson, F., Duperier, C., Garrier, W., Saunier, P.-E., Pitarch, Y., Jourdan, F., Giacomoni, F., Frainay, C.: FORUM: building a Knowledge Graph from public databases and scientific literature to extract associations between chemicals and diseases. Bioinformatics **37**(21), 3896–3904 (2021). doi:10.1093/bioinformatics/btab627

[7] Délos, S., Carsol, J.L., Ghazarossian, E., Raynaud, J.P., Martin, P.M.: Testosterone metabolism in primary cultures of human prostate epithelial cells and fibroblasts. The Journal of Steroid Biochemistry and Molecular Biology **55**(3-4), 375–383 (1995). doi:10.1016/0960-0760(95)00184-0
